# Sex-specific signatures of brain-wide induction of ΔFOSB and altered co-activation networks in a mouse model for exercise training

**DOI:** 10.64898/2026.02.24.707648

**Authors:** Marene H. Hardonk, Rick Wenning, Jazz Stofberg, Meike H. Mulder, Anna H. Vuuregge, Jorine Geertsema, Susanne E. la Fleur, Paul J. Lucassen, Joram D. Mul

## Abstract

Physical exercise training promotes brain health, yet the underlying mechanisms remain incompletely understood. Repeated neuronal activation results in accumulation of the transcription factor ΔFOSB, a long-lived splice variant of FOSB. We have previously demonstrated in rats that long-term voluntary wheel running (VWR), a behavioral paradigm that mimics exercise training in humans, altered ΔFOSB expression in a brain-wide manner, and that it reorganized co-activation networks. Here, we used a similar approach, now in mice, to map neuronal activation patterns following long-term VWR.

Young-adult male and female C57BL/6J^OlaHsd^ mice were allowed to run for four weeks on horizontal wheels, after which ΔFOSB immunoreactivity was quantified across 46 brain regions associated with stress regulation, cognition and reward-related behavior. Subsequently, network analysis was applied to assess VWR-mediated changes in patterns of interregional ΔFOSB co-activation and network topology.

Male and female mice ran equal distances and VWR blunted bodyweight gain and final fat mass in both sexes. VWR modulated ΔFOSB expression across several cortical, striatal, hippocampal and thalamic regions. Network analysis revealed a sex-specific network reorganization, with reduced overall network density and increased cortical centrality in males, and greater global efficiency (*i.e.* small-worldness) in females. Thus, VWR induced large-scale, sex-dependent adaptations in brain (in)activation, reshaping network organization in distinct ways across sexes.

Because ΔFOSB regulates many target genes, our findings indicate that long-term VWR induces widespread transcriptional alterations throughout the mouse brain. More focused follow-up studies are required to investigate the impact of these specific alterations on stress regulation, cognition and reward-related behavior.

**SIGNIFICANCE STATEMENT:** Exercise training promotes brain health, but the underlying mechanisms remain elusive. Here, we studied ΔFOSB, a very stable transcription factor involved in neuroplasticity, after four weeks of running in mice. We show this altered ΔFOSB expression in a subset of 46 brain regions implicated in stress regulation, cognition and reward, notably with sex-specific signatures.

This was accompanied by specific changes in ΔFOSB co-activation networks, including decreased network density and increased cortical centrality in males, and greater network efficiency in females.

Our mouse ΔFOSB brain map following running improves our understanding of how exercise training impacts brain plasticity and offers a framework for more mechanistic future studies into running-mediated changes in stress regulation, cognition and reward-related behavior.

## INTRODUCTION

Exercise training improves many physical health-related parameters, including several benefits for the brain. Associated with the latter are improvements in cognition, increases in stress resilience and a reduction in the incidence and severity of mental health disorders (Chekroud et al., 2018; Harvey et al., 2018; Kandola et al., 2016; Pearce et al., 2022; Schuch et al., 2016). So far, the exact neurobiological mechanisms through which exercise training promotes brain health remain elusive, but could provide valuable insights for therapeutic strategies based on exercise training to halt cognitive impairment or to increase stress resilience.

Animal experiments allow to map exercise training-induced adaptations at the whole-brain level that can possibly guide future mechanistic or therapeutic studies in humans. As such, voluntary wheel running (VWR), a self-reinforcing physical activity modality, is widely used in rodents to model aspects of human aerobic exercise training (Greenwood et al., 2011). VWR, similar to human exercise training (Chekroud et al., 2018; Harvey et al., 2018; Kandola et al., 2016; Pearce et al., 2022; Schuch et al., 2016), enhances cognitive performance, boosts stress resilience and reduces depressive-like behaviors in stress-exposed rodents (Donoso et al., 2023; Duman et al., 2008; Greenwood & Fleshner, 2019; Mul, 2018; Tanner et al., 2024).

A previously established molecular mechanism in rodents that mediates stress resilience during sedentary and VWR conditions is the induction of the transcription factor ΔFOSB in specific subpopulations of the nucleus accumbens (NAc), a central component of the brain’s reward system (Mul et al., 2018; Vialou, Robison, et al., 2010). ΔFOSB is a truncated, and highly stable, splice variant of the more classic form FOSB (Nestler, 2015), that forms activator protein-1 (AP-1) transcriptional complexes with JUN proteins, and that can, also via self-assemblies, regulate transcription in a broad set of downstream genes (Hope, 1998; McClung & Nestler, 2003; Yin et al., 2020). Due to its well-established long half-life (Carle et al., 2007; Chen et al., 1997; Ulery-Reynolds et al., 2009), ΔFOSB accumulates in neurons in particular following repetitive activation (Chen et al., 1997; Kelz & Nestler, 2000; McClung et al., 2004; Nestler et al., 2001) and is thus particularly suitable to map patterns of repeated neuronal activation across the whole brain, such as following long-term VWR.

We recently used this approach in male and female Wistar rats and reported widespread ΔFOSB expression across various stress-, reward-, and cognition-associated regions, as well as brain-wide changes in co-activation networks compared to sedentary control rats (Hardonk et al., 2026). Here, we extend this work to mice and quantified ΔFOSB expression across 46 stress-, cognition-, and reward-related brain regions in male and female C57BL/6J^OlaHsd^ mice following four weeks of VWR. Subsequently, we constructed ΔFOSB co-activation networks to assess how exercise training reorganizes brain-wide neuronal activity patterns and directly compared those networks between sexes to evaluate sex-specific effects. This mouse ΔFOSB brain atlas thereby provides a framework for more mechanistic studies into the changes that occur in neuronal activation following VWR, and into those related to e.g. neuronal excitability and stress-, cognition- and reward-related behaviors.

## METHODS AND MATERIALS

### Ethical approval and monitoring

All experimental procedures were performed in accordance with the European guidelines for laboratory animals (EU directive 2010\63\EU) and approved by the Dutch Central Committee for Animal Experiments (CCD; AVD8010020172424) and the Agency for Animal Welfare (IvD) of the Netherlands Institute of Neuroscience (NIN; Royal Dutch Academy of Sciences), Amsterdam, and the IvD of the University of Amsterdam, Amsterdam, as required by Dutch law.

### Animals and housing conditions

A graphical overview of the experimental procedures is presented in Fig. 1. We used 24 male (19-22g and approximately 6 weeks-of-age at arrival) and 24 female (18-22g and approximately 12 weeks-of-age at arrival) C57BL/6J^OlaHsd^ mice (strain code: 057, Inotiv, the Netherlands). Males and females were studied in two separate cohorts (*n* = 24/cohort). Mice were housed at the animal facility of the Swammerdam Institute for Life Sciences (SILS) at the University of Amsterdam in a temperature- (21-23°C), humidity- (40-60%) and light-controlled room (12:12h light/dark cycle) with lights on at Zeitgeber time 0 (ZT0; 08:00).

**Figure 1.**
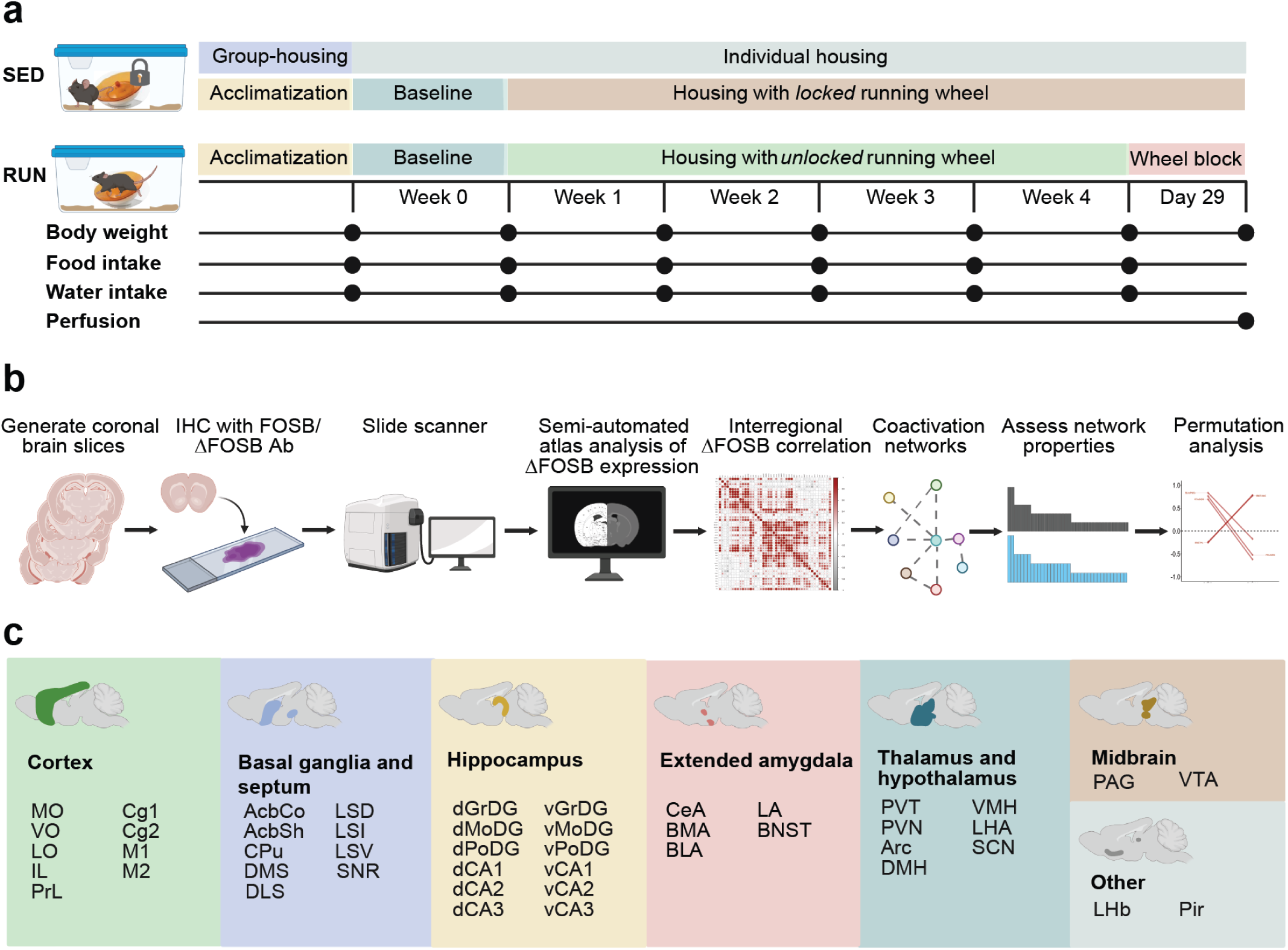
Experimental timeline and workflow. (**a**) Experimental timeline of animal studies. All male and female mice were housed with a locked running wheel (sedentary; SED) during the 7-day baseline period. During the subsequent 28-day experimental period, SED controls remained housed with a locked running wheel, whereas the running wheels of runners (RUN) were unlocked. Body weight, food and water intake were measured weekly. Wheels were blocked for 24h before isolation of brains. (**b**) Experimental workflow of immunohistochemistry, ΔFOSB quantification, and ΔFOSB co-activation network generation and analysis. (**c**) Brain regions of interest (ROIs) categorized by parent structures: cortex (green), basal ganglia and septum (purple), hippocampus (yellow), extended amygdala (pink), thalamus and hypothalamus (blue), midbrain (orange), and other (gray), see Supplementary Tables 2-1 and 3-1 for a list of abbreviations.

Mice were allowed to recover from transport stress and left undisturbed for seven days during group housing (4 animals/cage) in high-temperature polysulfane (PSU) type II long cages measuring 36.5 x 20.7 x 14.0 cm (length x width x height; 530cm², 1284L Eurostandard Type II L, Tecniplast) with standard wood-chip bedding and *ad libitum* access to a bottle of tap water and a pelleted diet [#801722, CRM (P), 22% protein, 69% carbohydrate, and 9% fat by energy, 3.6 kcal/g gross energy, Special Diet Services], unless mentioned otherwise, and with a wooden gnawing stick, and Enviro-Dri nesting material as cage enrichment. Continuous and soft radio sound provided background noise.

### Energy homeostasis and VWR

Following acclimatization to the SILS animal facility, RandoMice (Van Eenige et al., 2020) was used to randomly assign mice based on body weight to either one of the two groups: sedentary (SED; male, *n* = 12; female *n* = 12) and voluntary wheel running (RUN; male, *n* = 12; female, *n* = 12). Sample sizes were calculated to ensure power (> 0.8) and based on prior experience with this behavioral paradigm as well as previous experiments of comparable design. All mice were housed individually in PSU type II long individually ventilated cages (IVC; 39.6 x 21.5 x 17.2cm, 542cm², 1285L cage, Tecniplast) with the previously mentioned cage enrichment (a wooden gnawing stick and Enviro-Dri nesting material) as well as a red mouse hut (Bio Services) and a locked low-profile wireless inclined (horizontal) running wheel (15.5cm diameter, 10.16cm high, Med Associates Inc, ENV-047 and ENV-044 for RUN and SED groups, respectively) and *ad libitum* access to a bottle of tap water and the CRM (P) diet for a baseline period of seven days.

Following this baseline period, wheels were unlocked for only the RUN group for another 28 days. Wheel revolutions per cage were recorded per minute using the accompanying software provided by the manufacturer (Wheel Manager Data Acquisition Software, SOF-860, Med Associates Inc). Total distance run was calculated based on the total number of revolutions and the circumference traveled for each revolution [12.04cm/revolution, estimated from the external (15.5cm) and internal (9cm) diameter of the running wheels]. Throughout baseline and the VWR period, body weight, food intake and water intake were measured weekly. After 28d of running, the wheels of the RUN group were locked at the start of the light phase for 24h before sacrifice. This was done to allow the degradation of any residual full-length (regular) FOSB protein, such that all remaining immunoreactivity reflects only the accumulated ΔFOSB signal (Nestler et al., 2001; Perrotti et al., 2004).

Blinding during the experiment was not feasible due to the presence of an unlocked or locked running wheel; however, all data analyses were conducted by investigators who were blinded to the group identity. Two male runners were excluded from the experiment due to health-related complications of unknown origin: one animal was found dead in the home cage, and the other animal exhibited impaired health and a marked cessation of running behavior after one week and was therefore removed from the experiment. Data from these two mice was excluded from all analyses. One female runner was excluded from running wheel analyses and several ROIs of two sedentary animals were excluded from all ΔFOSB-related analyses due to technological failures (*i.e.* failed running wheel recordings and unsuccessful ΔFOSB staining, respectively).

### Tissue collection and single-labeling immunohistochemistry

After the 24h wheel blockade, mice were fasted starting at ZT0 and sacrificed between ZT0 and ZT8 via an overdose of intraperitoneally administered pentobarbital(Euthasol; 120mg/kg). Gonadal white adipose tissue (gWAT) pads and adrenals were bilaterally removed and weighed prior to transcardial perfusion with ice-cold 0.9% saline (0.9% w/v NaCl in ultrapure H_2_O) followed by ice-cold 4% paraformaldehyde (PFA) in 0.1M phosphate buffer (PB; pH 7.4). Afterwards, brains were dissected and postfixed in 4% PFA in PBS for 24 hours at 4°C, then washed in PBS and transferred to a 0.01% w/v sodium azide in 0.1M PB solution and stored in 4°C until use.

Prior to sectioning, brains were cryoprotected in 15% (w/v) sucrose in 0.1M PB at 4°C until saturated (*i.e* completely sunken to the bottom of the falcon tube), followed by overnight immersion in 30% (w/v) sucrose in 0.1M PB at 4°C until saturation. Coronal brain sections (35μm) were cut using a sliding microtome (Jung HN40, Leica Biosystems) and stored in cryoprotectant solution (20% v/v glycerol, 30% v/v ethylene glycol in 0.05M PBS) at −20°C. Sections containing the cortex, basal ganglia/septum, hippocampus, (extended) amygdala, thalamus, hypothalamus and midbrain were collected from Bregma +2.80mm to Bregma −4.04mm (Paxinos & Franklin, 2001).

Visualization of ΔFOSB signal was performed as previously described (Hardonk et al., 2026). In short, free-floating sections from male brains were washed in 0.05M tris-buffered saline (TBS), endogenous peroxidase activity was inhibited by pretreating for 10 minutes with 1.5% v/v H_2_0_2_ (Merck; in TBS) before incubation with rabbit anti-FOSB [5G4; #2251, Cell Signaling Technology, RRID: AB_2106903; 1:1000 in supermix (0.25% w/v gelatin, 0.5% v/v Triton X-100, in 1x TBS, pH 7.6)] for 1h at room temperature (RT) and overnight at 4 °C. Following incubation with a biotinylated goat anti-rabbit IgG (H + L) (BA-1000; Vector Laboratories, Burlingham, RRID: AB_2313606; 1:200 in supermix) for 2h at RT, signal amplification was achieved using Avidin-Biotin complex (Vectastain Elite ABC HRP Kit, PK-6100, Vector Laboratories; 1:800 in supermix) for 1.5h at RT, and visualized by nickel-enhanced DAB [0.05% w/v 3.3 -diaminobenzidine (Sigma), 0.23% w/v nickel-ammoniumsulphate (Merck) and 0.01% w/v H_2_O_2_ (Merck) in 0.05M TB, pH 7.6].

Sections were then mounted onto Superfrost Plus slides (Thermo Scientific), air-dried, dehydrated through graded ethanol and cleared in Xylene, before slides were coverslipped with Entellan (#107961, Merck). For female brain sections, antigen retrieval was necessary to facilitate quantifiable ΔFOSB staining comparable to that observed in males. Sections were then mounted on SuperFrost Plus slides, outlined with a PAP-PEN liquid blocker and dried overnight. Slides were then washed in TBS and antigen retrieval was performed by microwaving the slides in 0.05M TB (pH = 9.0) for 15 minutes (800 Watt, heat reduced to 240 Watt when boiling). After cooling for 30 min, ΔFOSB immunohistochemistry was performed as described above.

### Image acquisition and analysis

For quantification of ΔFOSB-positive nuclei and construction of co-activation networks, an experimentalpipeline was used similar to that previously described in Hardonk et al. (2026). In short, randomly numbered coronal brain sections were scanned at 10X Brightfield magnification using a Zeiss Axio Scan Z1 slide scanner (Carl Zeiss AG, Oberkochen, Germany) to ensure blindness to group identity during analysis. ΔFOSB-expressing cells per region of interest (ROI) were quantified usinga modified version of the atlas-based analysis (Bourgeois et al., 2021) in FIJI (ImageJ2), which was adapted in-house for bilateral mouse brain sections. Sections were converted to 8-bit grayscale, aligned to a mouse brain atlas (Bourgeois et al., 2021) and thresholded (Robust Automatic Threshold Selection plugin [noise = 25, lambda = 3]; Analyze Particle function [size = 20 - 250, circularity = 0.40 - 1.00]) to detect strongly labeled ΔFOSB-positive nuclei. Counts were normalized to ROI area and mean of the respective sedentary control group. Missing or damaged ROIs were excluded. An overview of included ROIs and full regional details are provided in Fig. 1C and Supplementary Tables 3-1 and 4-1.

### Co-activation network generation and permutation analyses

Co-activation network analyses were conducted as previously described ((Hardonk et al., 2026; Terstege & Epp, 2022). Pearson correlations between regional ΔFOSB expression levels were calculated in R (version 4.4.1) to generate group-specific correlation matrices and were visualized as unweighted (*i.e.* unthresholded) and weighted (*i.e.* thresholded) heatmaps. From these matrices, co-activation networks were constructed by thresholding correlations at P < 0.005 using the igraph package (R version 2.1.4), with brain regions represented as nodes and significant correlations as edges.

To further describe the co-activation networks, global (*i.e.* density, small-worldness) and local (*i.e.* degree, weighted betweenness centrality) network metrics were computed using the igraph package following established methods (Bullmore & Bassett, 2011; Newman, 2003; Onnela et al., 2005; Rubinov & Sporns, 2010; Watts & Strogatz, 1998). To assess whether VWR or sex reorganized the overall network architecture, similarity between the two correlation matrices of interest was evaluated using a Mantel test with 999 permutations. To identify which specific interregional correlations were altered, correlations from one group were subtracted from the other and group differences in correlation coefficients were evaluated using permutation testing (*n* =1000) (Jin et al., 2024).

### Statistical analysis

All behavioral, metabolic, and physiological data were analyzed using two-tailed unpaired Student’s t-tests, or Mann–Whitney U tests when parametric assumptions were violated, and two-way ANOVAs with repeated measures where appropriate, followed by Šidák post hoc tests. Regional ΔFOSB analyses included false discovery rate (FDR) correction for multiple comparisons using the Benjamini–Hochberg procedure, with an FDR threshold of *P*_corrected_ < 0.05, and both uncorrected and FDR-corrected *P* values are reported. Correlations were assessed using simple linear regression. For all behavioral, metabolic and physiological data, as well as regional ΔFOSB analyses, statistical analyses and data visualization was conducted in GraphPad Prism (v10.2.3) and *P* < 0.05 was considered significant, while *P* < 0.10 was considered a trend towards an effect. See figure legends and Supplementary Tables 3-1 and 4-1 for full statistical details.

Interregional ΔFOSB expression correlations were computed and visualized in unfiltered heatmaps. Networks were constructed and analyzed based on thresholded correlation matrices (*P* < 0.005). Overall matrix differences between groups were assessed using a Mantel test with 999 permutations, with *P* < 0.05 considered statistically significant. Permutation testing of individual correlation differences between experimental groups was performed using bootstrapping (n = 1000), with unweighted and significant results (*P* < 0.005) visualized in heatmaps and volcano plots (with additional threshold for correlation differences of >1). All network analyses, statistical computations and visualizations were performed in R (version 4.4.1).

## RESULTS

### Long-term VWR behavior

To examine how long-term VWR influenced ΔFOSB expression and brain-wide co-activation networks, young adult male and female mice were housed individually with either a locked running wheel (controls) or with an unlocked wheel for 28 days (runners; Fig. 1A). Both sexes gradually increased their daily running distances over the first days, after which running activity stabilized at approximately 13 km/day in males and 17 km/day in females (Fig. 2A, B). Across the 28-day period, males and females ran a total distance of 351.2 ± 38.3 km and 432.5 ± 24.9 km on average, respectively, with no significant differences between sexes (Fig. 2C). As expected with a nocturnal species, most running occurred during the dark phase (Supplementary Fig. 2-1A–J). Overall, both males and females displayed robust habitual VWR.

**Figure 2.**
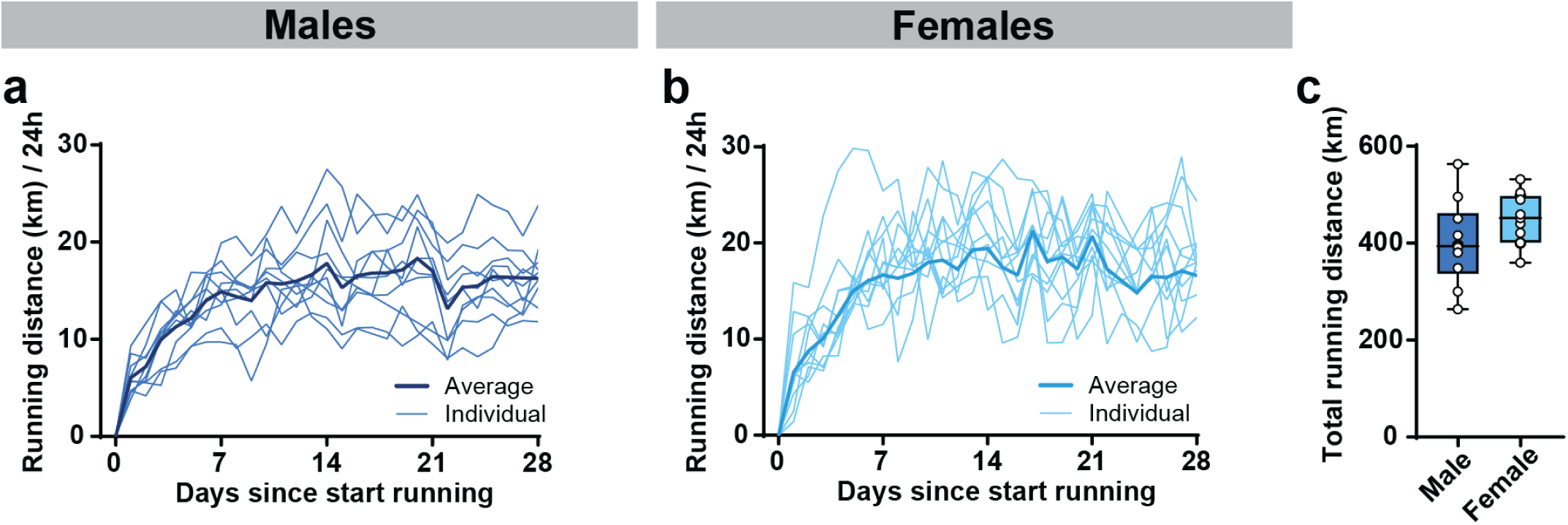
Running behavior in males and females. (**a, b**) Daily individual and average running distances of (a) males and (b) females. (**c**) Total running distance of males and females during 28 days of running (t_(19)_ = 1.593, *P* = 0.128). Data are presented as the mean and individual lines representing individual mice (a, b) as box plots indicating the median (line) and mean (+), the interquartile range, and the minimum to maximum values of the data distribution, with dots representing individual mice (c). (a-c) *n* = 10-11/group.

### Physiological and metabolic impact of long-term VWR

Both male and female mice, irrespective of running condition, gained body weight during the experimental period. However, male runners exhibited significantly lower body weight gain compared to sedentary controls from week 2 onwards, whereas no significant differences were detected between female runners and sedentary controls during the entire VWR period (Supplementary Fig. 2-2A, C). Thus, VWR attenuated body weight gain exclusively in males (Supplementary Fig. 2-2A–D).

Running reduced daily caloric intake compared to sedentary controls during the first week in males (Supplementary Fig. 2-2E). During weeks 2–4, male and female runners consumed significantly more calories than their sedentary counterparts (Supplementary Fig. 2-2E, G). After 28 days, cumulative caloric intake did not differ between male runners and controls but was higher in female runners compared to female controls (Supplementary Fig. 2-2F, H).

Running also impacted daily water consumption (Supplementary Fig. 2-2I–L). From week 2 onwards, both male and female runners showed a trend to consume more water than their respective sedentary counterparts; however, post-hoc analyses did not reveal significant group differences, and cumulative water intake did not differ between conditions (Supplementary Fig. 2-2I–L).

After 28 days, terminal gWAT weight was lower in both male and female runners compared to their respective sedentary controls (Supplementary Fig. 2-3A, C). Moreover, terminal gWAT weight negatively correlated with total running distance in males, whereas no correlation was observed in females (Supplementary Fig. 2-3B, D). In contrast, adrenal weight normalized to body weight was not significantly affected by running in either sex, nor were correlations observed with total running distance (Supplementary Fig. 2-3E–H).

**Figure 3.**
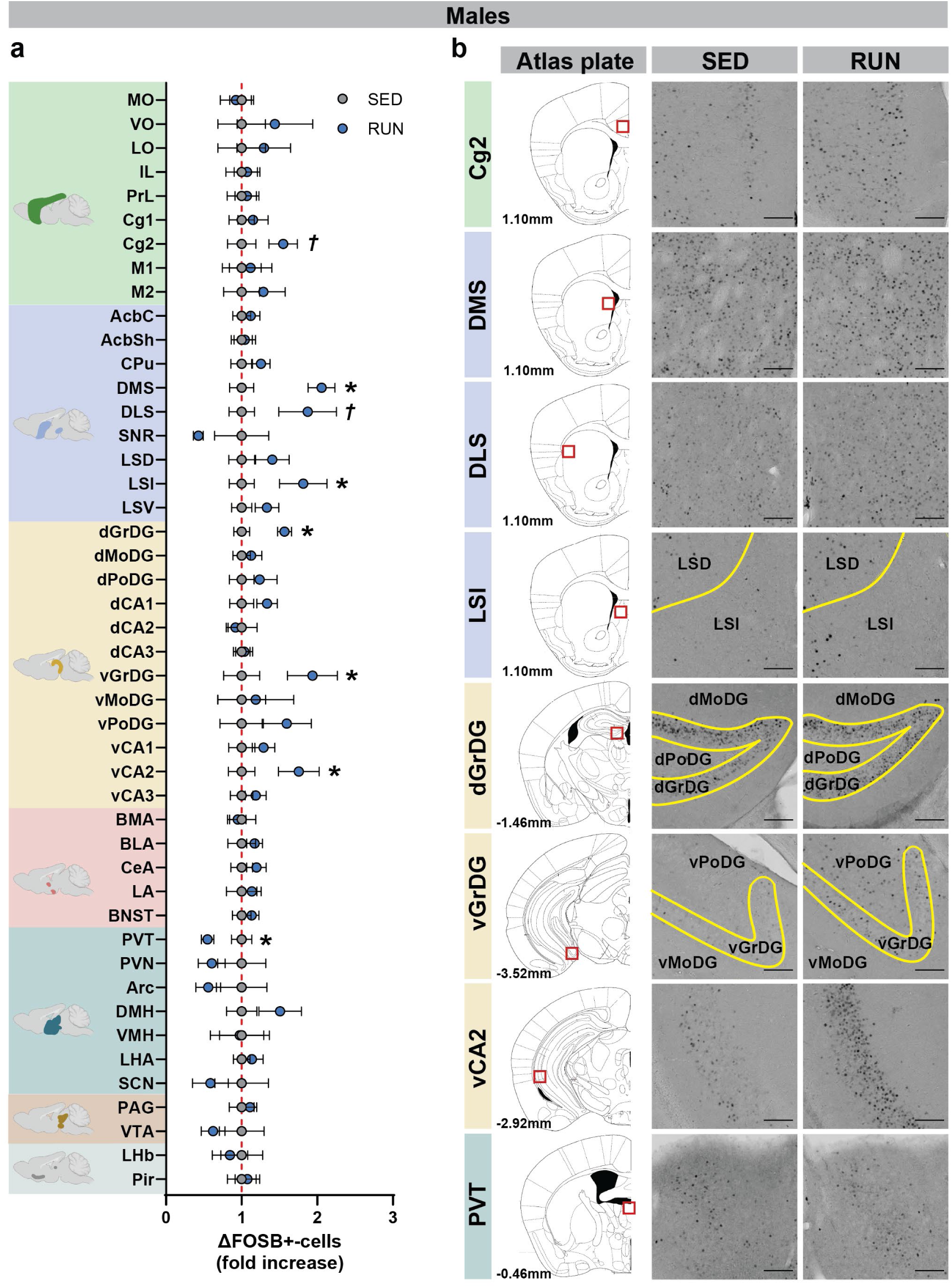
Long-term VWR induction of ΔFOSB in males. (**a**) Fold change in the mean number of ΔFOSB-positive cells (per mm²) per region of interest (ROI) in male sedentary (SED) controls and runners (RUN), normalized to the mean of SED mice; \**P* < 0.05, *†P* < 0.10 before FDR correction; see Supplementary Table 3-1 for exact statistical values. (**b**) Representative images of ΔFOSB immunoreactivity in the cingulate cortex 2 (Cg2), dorsomedial striatum (DMS), dorsolateral striatum (DLS), intermediate part of the lateral septum (LSI), the dorsal (dGrDG) and ventral (vGrDG) part of the granule cell layer of the dentate gyrus, the ventral cornu ammonis 2 (vCA1) and the paraventricular thalamic nucleus (PVT). Scale bar = 100 µm. Data are presented as the mean ± S.E.M (a). (a) *n* = 10-12/group.

**Figure 4.**
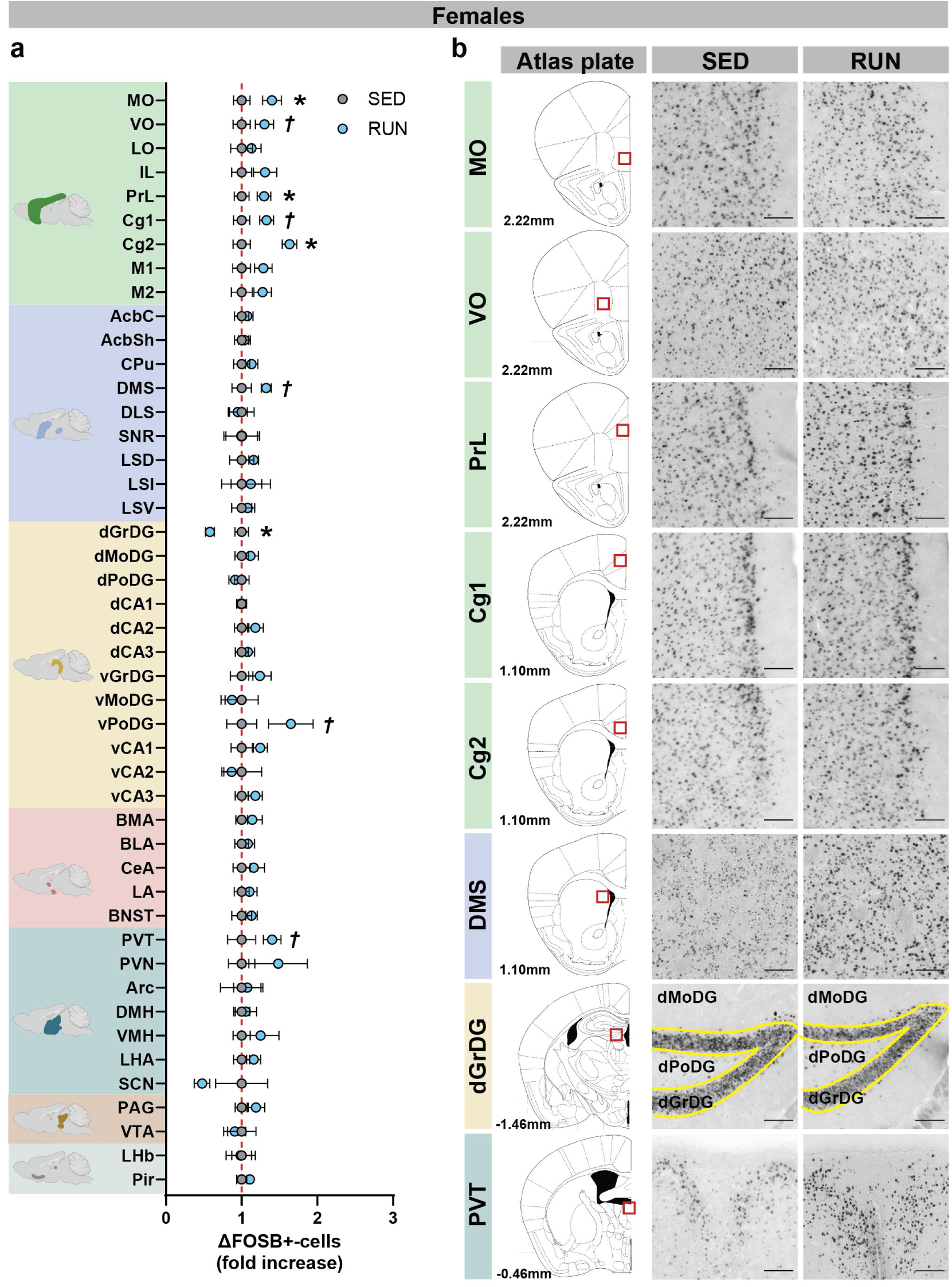
Long-term VWR induction of ΔFOSB in females. (**a**) Fold change in the mean number of ΔFOSB-positive cells (per mm²) per region of interest (ROI) in female sedentary (SED) controls and runners (RUN), normalized to the mean of SED mice; \**P* < 0.05, *†P* < 0.10 before FDR correction; see Supplemental Table 4-1 for exact statistical values. (**b**) Representative images of ΔFOSB immunoreactivity in the medial (MO) and ventral orbital cortex (VO), prelimbic cortex (PrL), cingulate cortex 1 (Cg1) and 2 (Cg2), dorsomedial striatum (DMS), dorsal part of the granule cell layer of the dentate gyrus (dGrDG) and the paraventricular thalamic nucleus (PVT). Scale bar = 100 µm. Data are presented as the mean ± S.E.M (a). (a) *n* = 12/group.

In summary, long-term VWR limited body weight gain and lowered adiposity, particularly in males, without significantly altering adrenal weight.

### Long-term VWR modulates regional ΔFOSB in males and females

Next, we evaluated the impact of long-term VWR on ΔFOSB-positive cell numbers in 46 brain regions associated with stress-, cognition- and reward-related behavior in male and female mice. To do this, we adapted an atlas-based analysis workflow (Bourgeois et al., 2021) for bilateral mouse brain sections and compared these outcomes with those of sedentary controls (Fig. 1B, C; see Supplementary Table 3-1 and 4-1 for a full overview of all included ROIs, Bregma ranges, number of slices included per group, percentage of change in runners and statistics).

In males, running significantly increased ΔFOSB-positive cell numbers in the dorsomedial striatum (DMS), the intermediate part of the lateral septum (LSI), the dorsal (dGrDG) and ventral (vGrDG) granule cell layers of the dentate gyrus, and the ventral cornu ammonis 2 (vCA2) (all *P* < 0.05; Fig. 3A, B; Supplementary Table 3-1). Male runners also showed a trend towards increased ΔFOSB-positive numbers in the cingulate cortex area 2 (Cg2) and the dorsolateral striatum (DLS) (both *P* < 0.10; Fig. 3A, B; Supplementary Table 3-1).

In contrast, running significantly decreased ΔFOSB-positive cell numbers in the paraventricular thalamic nucleus (PVT) (*P* < 0.05; Fig. 3A, B; Supplementary Table 3-1). After applying FDR correction for multiple comparisons, only the increases observed in the DMS and dGrDG remained statistically significant (Supplementary Table 3-1). A trend towards a negative correlation between total running distance and ΔFOSB-positive numbers was observed in the ventral molecular layer of the dentate gyrus (vMoDG) (Supplementary Fig. 3-1A, B), but this correlation did not survive multiple comparisons correction.

In females, running significantly increased ΔFOSB-positive cell numbers in the medial orbital cortex (MO), Cg2, and prelimbic cortex (PrL; *P* < 0.05), with trends towards an increase in the ventral orbital cortex (VO), Cg1, DMS, ventral polymorph layer of the dentate gyrus (vPoDG) and PVT (all *P* < 0.10). Conversely, running significantly reduced ΔFOSB-positive cell numbers in the dGrDG. After FDR correction, only the effects in the DMS and dGrDG remained statistically significant (Supplementary Table 4-1). Running distance correlated positively with ΔFOSB-positive cell numbers in the dorsal molecular layer of the dentate gyrus (dMoDG) and vCA3, and showed a trend for a positive correlationin the lateral habenula (LHb; Supplementary Fig. 4-1A, E-G). Running distance negatively correlated with ΔFOSB-positive cell numbers in the dorsal part of the lateral septum, and showed a trend (*P* < 0.10) for negative correlations with ΔFOSB-positive cell numbers in the MO and VO (Supplementary Fig. 4-1A-D). However, none of these (significant) correlations survived FDR correction for multiple comparisons.

Taken together, these observations demonstrate that long-term VWR impacts ΔFOSB expression in several brain regions involved in stress-, cognition- and reward-related behavior in males and females.

### Long-term VWR alters brain ΔFOSB co-activation network efficiency and topology in males

To further explore how exercise influenced ΔFOSB expression across stress-, cognition-, and reward-related circuits at the network level, we analyzed ΔFOSB co-activation patterns by cross-correlating regional ΔFOSB expression in male and female sedentary controls and runners. For further network characterization, we quantified global (*i.e.* density, small-worldness) and local (*i.e.* degree, betweenness centrality) network metrics.

Sedentary males exhibited a densely interconnected network, as shown by both the co-activation heatmap and network visualizations, with multiple regions across distinct structures occupying central positions(Fig. 5A, B). Consistent with this pattern, network density was relatively high in sedentary controls (*k_DEN_* = 0.278; Fig. 5C). By contrast, male runners showed an overall modest reduction in connectivity (*k_DEN_* = 0.251; Fig. 5C), accompanied by a pronounced shift towards greater small-world-like organization (SED: *σ* = 2.294; RUN: *σ* = 2.793; Fig. 5D). As a higher small-worldness coefficient reflects greater clustering (transivity) and shorter average path lengths, this observed increase in small world-like properties suggests enhanced global network efficiency. These group differences were robust across multiple correlation thresholds (α = 0.05, 0.01, 0.0001; Supplementary Fig. 5-1A, B), confirming that VWR reliably alters network organization in males.

**Figure 5.**
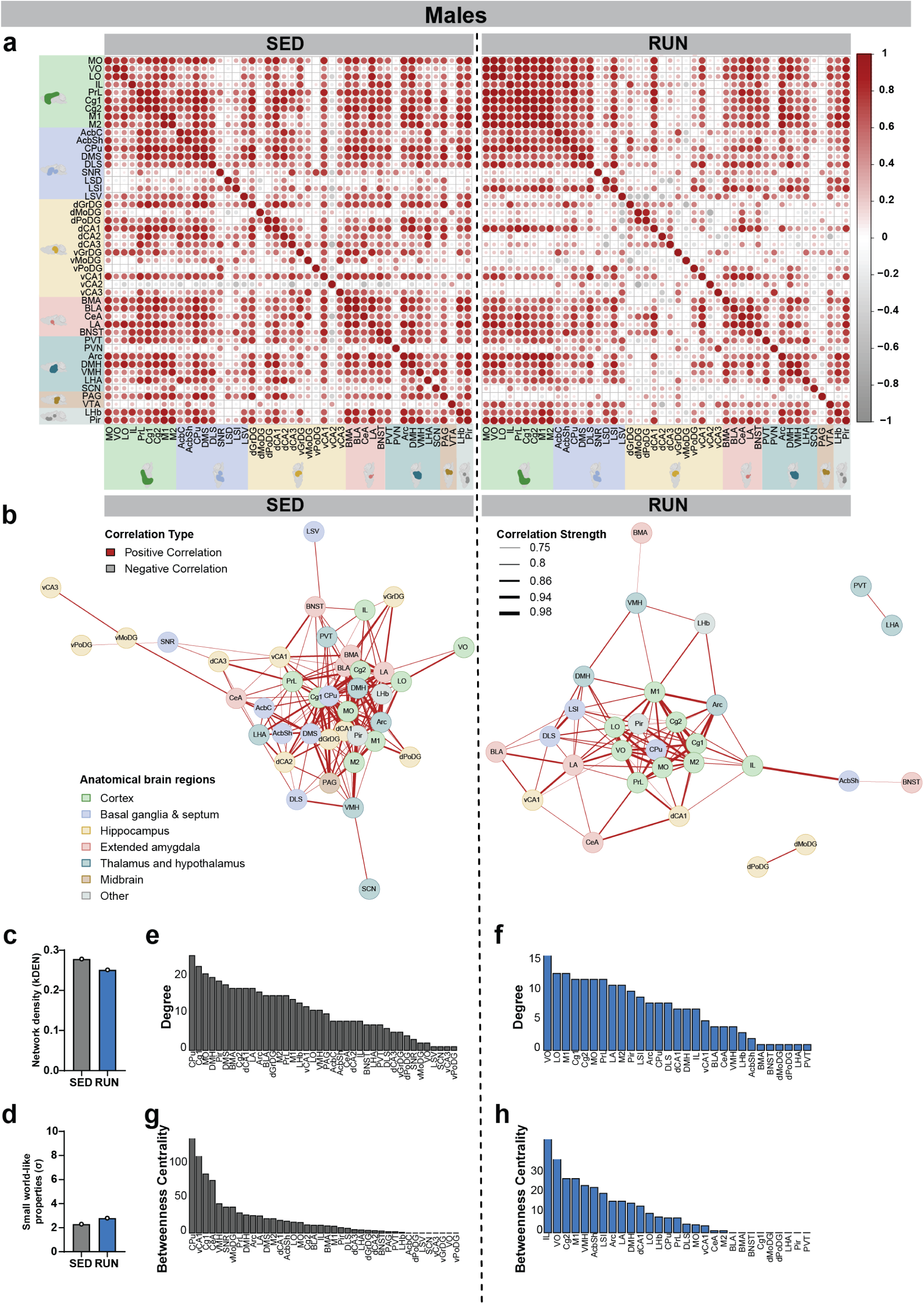
Network analysis reveals altered ΔFOSB co-activation following VWR in males. (**a**) Unweighted regional cross-correlation matrices of ΔFOSB-positive cells per mm² between all pairs of neuroanatomical regions of male sedentary controls (SED) and runners (RUN). (**b**) ΔFOSB co-activation networks constructed after thresholding for the strongest and most significant correlated or anti-correlated connections (r > 0.7, P < 0.005) in SED and RUN mice. (**c**) Network density expressed as *k_DEN_* of SED and RUN mice. (**d**) Small world-like properties expressed as *σ* of SED and RUN mice (**e, f**) Node degree per ROI of (e) sedentary controls (SED) and (f) male runners (RUN). (**g, h**) Betweenness centrality per ROI of (g) SED and (h) RUN mice. (a-h) *n* = 10-12/group. (b-h) *P* < 0.005 was considered significant.

Local network metrics further provided insight into the structural shift observed at the whole-network level. Both male runners and controls showed a long-tailed distribution of degree and betweenness centrality, reflecting the presence of strongly influential hub nodes (Fig. 5E-H). In sedentary males, high-degree nodes were predominantly located in the striatum [caudate putamen (CPu)], cortex (Cg1, MO), hypothalamus [dorsomedial hypothalamus (DMH)], and piriform cortex (Pir), consistent with the broad, multisystem centrality observed in the network visualizations (Fig. 5B, E). Betweenness centrality analysis supported this architecture, identifying striatal (CPu), hippocampal (vCA1), cortical (Cg1), amygdalar (CeA), and hypothalamic [ventromedial hypothalamus (VMH)] as dominant hubs (Fig. 5G). In male runners, both degree and betweenness shifted prominently towards cortical regions—including VO, LO, M1, Cg1, Cg2, MO, PrL, and infralimbic cortex (IL)—reflecting the cortico-centric architecture also identified in the co-activation maps (Fig. 5F, H).

Taken together, these results demonstrate that long-term VWR fundamentally reshapes ΔFOSB co-activation networks in males by lowering overall connectivity, improving network efficiency, and reallocating hub status towards cortical systems. This suggests a reorganization that favors a more streamlined, cortical-centered information flow, while reducing diffuse connectivity.

### Long-term VWR alters ΔFOSB co-activation network efficiency and topology in females

In sedentary females, co-activation heatmaps and network plots revealed two smaller networks: one primarily involving the prefrontal cortex, dorsal striatum and Pir, and the other primarily involving the hippocampus, amygdala and lateral hypothalamus, interconnected via the bed nucleus of the stria terminalis (BNST; Fig. 6A, B). The presence of these two partially segregated subnetworks is relevant for the interpretation of local network metrics, as measures such as degree and betweenness centrality may be distributed across subnetworks rather than concentrated within a single integrated network.

**Figure 6.**
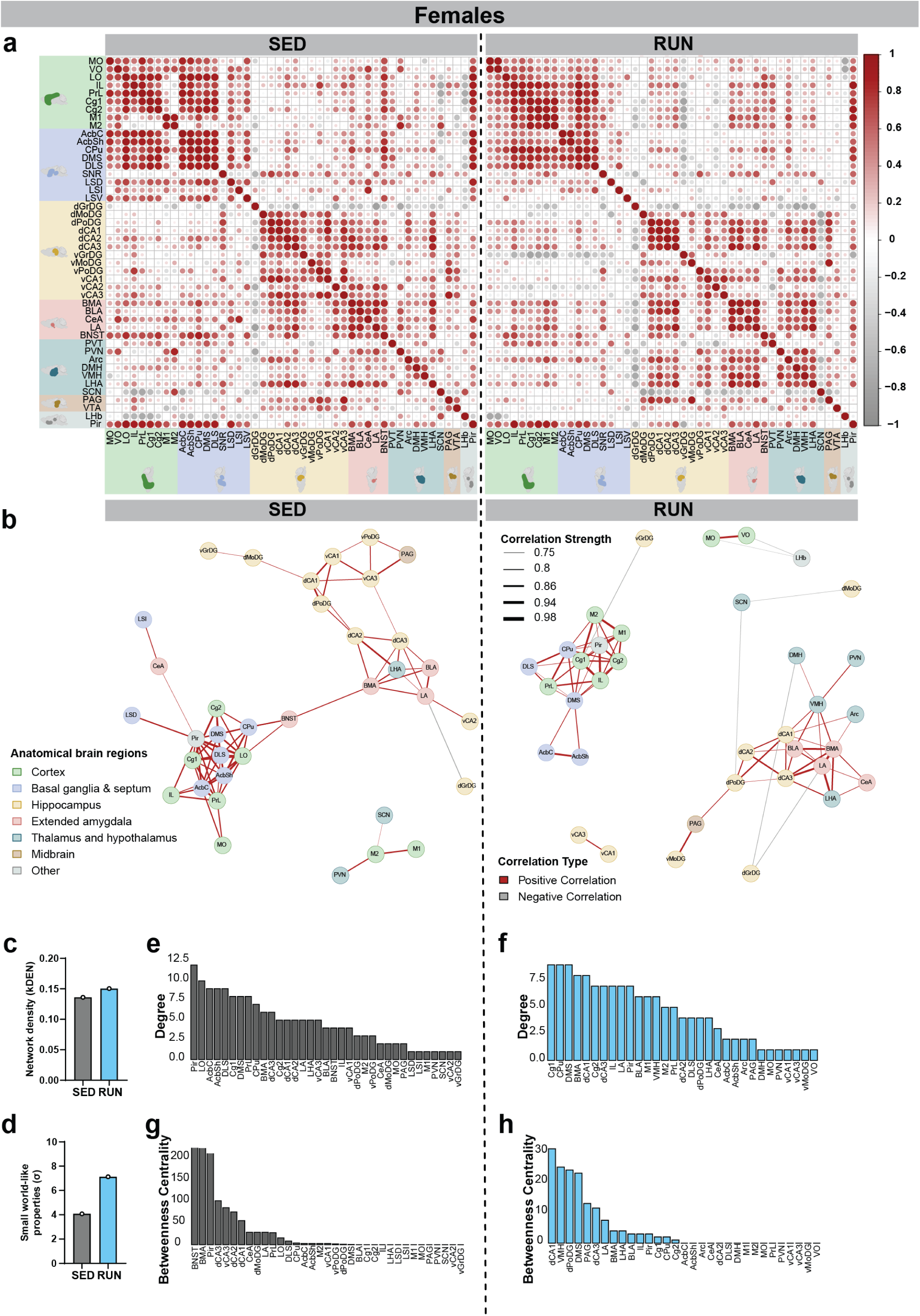
Network analysis reveals altered ΔFOSB co-activation following VWR in females. (**a**) Unweighted regional cross-correlation matrices of ΔFOSB-positive cells per mm² between all pairs of neuroanatomical regions of female sedentary controls (SED) and runners (RUN). (**b**) ΔFOSB co-activation networks constructed after thresholding for the strongest and most significant correlated or anti-correlated connections (r > 0.7, P < 0.005) in SED and RUN mice. (**c**) Network density expressed as *k_DEN_* of SED and RUN mice. (**d**) Small world-like properties expressed as *σ* of SED and RUN mice (**e, f**) Node degree per ROI of (e) sedentary controls (SED) and (f) female runners (RUN). (**g, h**) Betweenness centrality per ROI of (g) SED and (h) RUN mice. (a-h) *n* = 10-12/group. (b-h) *P* < 0.005 was considered significant.

In female runners, co-activation networks largely resembled those of their sedentary counterparts but showed subtle differences, including greater involvement of motor areas and reduced centrality of orbital cortex, nucleus accumbens and ventral hippocampal regions (Fig. 6B). Additionally, dorsal hippocampal and amygdala regions exhibited stronger connections with hypothalamic areas but weaker connections with the lateral hypothalamus (Fig. 6B). Network density was lower in sedentary females compared to sedentary males (*k_DEN_* = 0.136; Fig. 6C), whereas female runners showed a modest increase in connectivity (*k_DEN_* = 0.151; Fig. 6C). These changes were accompanied by a marked increase in small-world-ness (SED: *σ* = 4.075; RUN: *σ* = 7.127; Fig. 6D) and were consistent across multiple thresholds (α = 0.05, 0.01, 0.0001; see Supplementary Figs. 6-1A, B). At the most stringent threshold (*P* < 0.0001), connections were largely confined to cortical regions, indicating that within-cortex connectivity represented the strongest network interactions in female runners (See Supplementary Figs. 6-1A, B).

Local network metrics further highlighted subtle structural shifts. Both running and sedentary females displayed a long-tailed distribution of degree and betweenness centrality, reflecting strongly influential hub nodes (Fig. 6E-H). In sedentary females, high-degree nodes were primarily cortical (LO), striatal (AcbC, AcbSh, DLS) or the Pir (Fig. 6B, E), aligning with one of the subnetworks observed (Fig. 6A, B). Remarkably, betweenness centrality analysis identified hubs corresponding to the second network, including (extended) amygdalar [BNST, basomedial amygdala (BMA)] and hippocampal (dCA3, vCA3, dCA2, dCA1) areas (Fig. 6A, B, G). In female runners, degree and betweenness shifted slightly, but still highlighted striatal (CPu, DMS), amygdalar (BMA) and hippocampal (dCA1, and dPoDG) regions as main hubs, while the ventromedial hypothalamus (VMH) and periaqueductal gray (PAG) emerged as new central nodes (Fig. 6F, H).

Taken together, these results demonstrate that also in females VWR fundamentally reshapes ΔFOSB co-activation networks by increasing global efficiency, while generally maintaining network topology.

### Permutation analyses reveal specific connections and broad connection patterns impacted by long-term VWR and sex

Next, we applied permutation testing to compare correlation matrices between experimental groups and to identify individual edges showing significant changes between groups. To visualize overall patterns of change, we first computed an unthresholded difference matrix by subtracting correlation coefficients of sedentary males from those of running males and displayed these values as a heatmap (Fig. 7A, left). This revealed stronger correlations among cortical regions, as well as between cortical regions with other brain regions, such as striatal regions and ventral hippocampal regions, in male runners (Fig. 7A, left). Furthermore, we observed a pattern of reduced correlations between mainly dorsal hippocampal regions and other brain regions, such as cortical regions and striatal regions (Fig. 7A, left). In addition, several correlations with amygdala, ventral hippocampal, (hypo)thalamic regions, as well as with the VTA, were stronger in male runners (Fig. 7A, left).

**Fig. 7:**
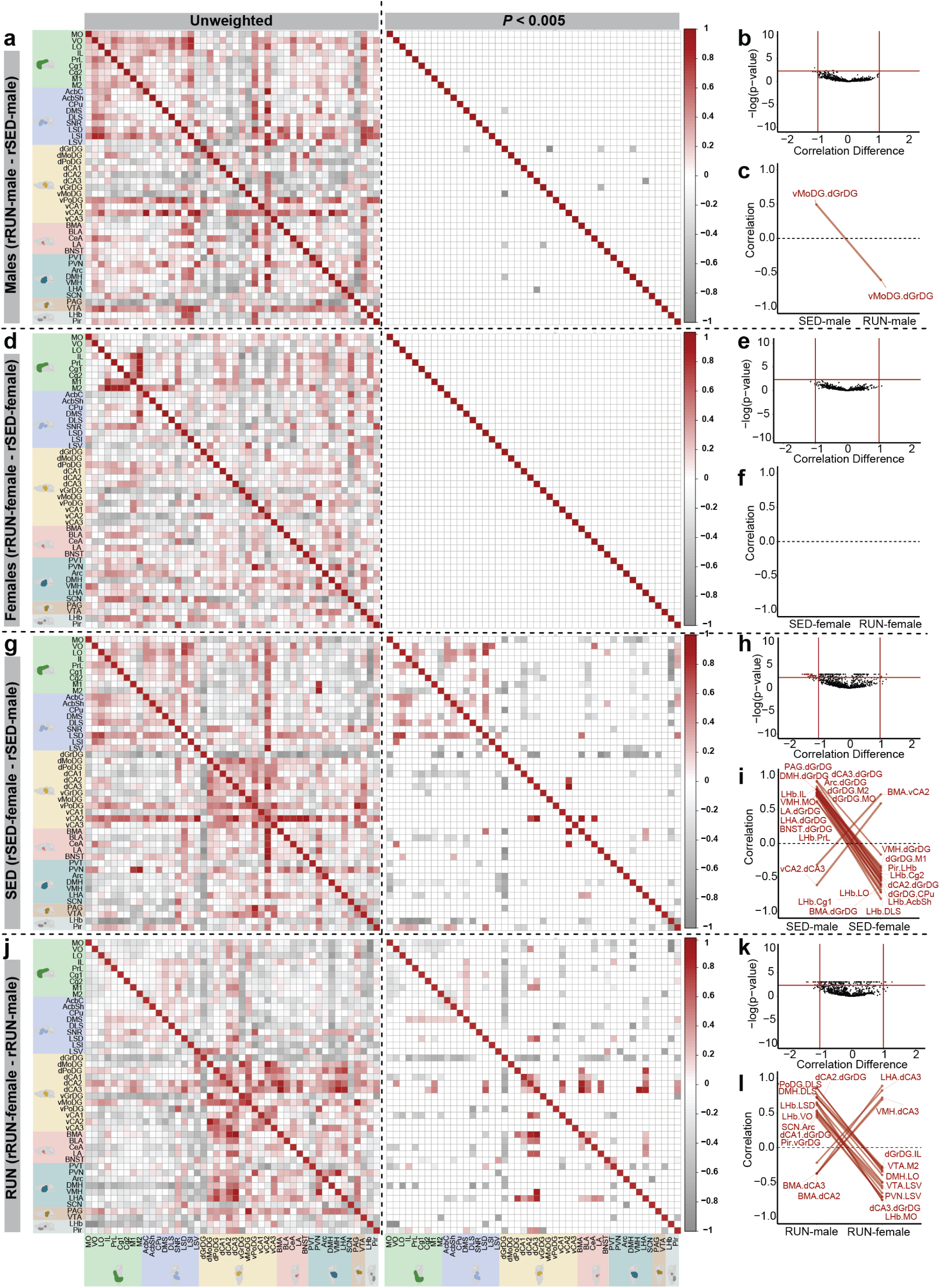
Permutation analysis reveals sex-specific long-term VWR-mediated alterations in ΔFOSB co-activation networks. **(a, d, g, j)** Unweighted and weighted (*P* < 0.005) heatmaps of Pearson correlation differences (r_group2_ – r_group1_) for all individual regional connections calculated from a permutation. The Mantel test revealed significant overall differences between the unweighted heatmaps of male and female sedentary (SED) and runners (RUN) (males: Mantel’s z = 225.168, *P* = 0.001, Fig. 7A; females: Mantel’s z = 141.206, *P* = 0.001; Fig. 7D) as well as between male and female runners (Mantel’s z = 198.995, *P* = 0.001; Fig. 7J), but not male and female sedentary controls Mantel’s z = 129.700, *P* = 0.135, Fig. 7G). (**b, e, h, k**) Volcano plot of Pearson correlation differences (r_group2_ – r_group1_) for all individual regional connections against their P-values calculated from a permutation analysis. Points intersecting or within the upper left or right quadrant represent the regional relationships with the greatest change (|correlation difference| > 1) that were most significant. (**c, f, I, l**) Parallel coordinate plots highlighting individual significantly changed regional correlations between groups. (a-l) n = 9-12/group. (a-l) *P* < 0.005 was considered significant.

To determine whether these changes reflected a reorganization of global network structure, we compared to full co-activation matrices using a Mantel test, which revealed a significant difference in overall co-activation matrices between male runners and sedentary controls (Fig. 7A, left). Next, to reveal local changes in the network, we performed permutation testing on individual correlations using a stringent threshold (*P* < 0.005) and identified significantly reduced correlations involving several hippocampal nodes (i.e. dGrDG-DMS, dGrDG-vMoDG, dGrDG-LA, dGrDG-DMH, dCA3-LHA, and vGrDG-LA; Fig. 7A, right). To detect correlations that reversed direction between groups, we applied an additional criterion [*P* < 0.005; r(difference) > 1], which revealed that VWR produced a shift in the relationship between the ventral molecular layer of the dentate gyrus (vMoDG) and the dGrDG, converting a positive correlation in male sedentary controls into a negative correlation in male runners (Fig. 7B, C). Collectively, these findings indicate that VWR in males produces both global and local shift in the co-activation networks of male mice.

In females, visualization of unthresholded difference matrices revealed more spatially dispersed alterations in correlations between sedentary and running mice (Fig. 7D, left). Despite this distributed pattern, comparison of the full co-activation matrices using a Mantel test indicated a significant difference in overall co-activation structure between female runners and sedentary controls (Fig. 7D, left). However, permutation testing of individual interregional correlations using the same stringent criteria [P < 0.005; with or without r(difference) > 1] did not identify any specific connections that differed significantly between groups (Fig. 7D, right; Fig. 7E, F), suggesting that VWR-related changes in females were broadly distributed across the network rather than driven by large shifts in specific correlations.

To evaluate baseline sex differences, we visualized unthresholded correlation differences by subtracting values of sedentary males from those of sedentary females (Fig. 7G, left). This comparison revealed stronger correlations in sedentary females among cortical, striatal, ventral hippocampal, and amygdala regions, accompanied by reduced correlations involving dorsal hippocampal regions. To determine whether these changes reflected differences in global network organization, we compared full co-activation matrices using a Mantel test. Despite region-specific alterations, overall interregional correlation structure did not differ significantly between sedentary males and females (Fig. 7G, left).

To identify specific connections underlying these local differences, permutation testing was applied using stringent criteria [*P* < 0.005; r(difference) > 1]. The most pronounced downregulated connections in sedentary females involved the dGrDG, which showed reduced correlations with amygdala (BNST, BMA, lateral amygdala [LA]), hypothalamic (LHA, DMH, VMH, arcuate nucleus [Arc]), (motor) cortical (MO, M1, M2), hippocampal (dCA2, dCA3), striatal (CPu), and midbrain (PAG) regions (Fig. 7H, I). In addition, LHb connectivity with cortical (LO, PrL, IL, Cg2, Cg1), striatal (AcbSh, DLS), and Pir areas was reduced, as was connectivity between the VMH and MO. Conversely, correlations involving the vCA2 with dCA3 and BMA were increased in sedentary females compared with sedentary males (Fig. 7H, I). Together, these findings indicate localized sex differences in specific interregional correlations at baseline without a corresponding shift in global network architecture.

Finally, we examined sex differences following VWR. Visualization of unthresholded difference matrices comparing male and female runners revealed stronger hippocampal connectivity in females, excluding connections involving dGrDG and dMoDG. Enhanced correlations were observed both within hippocampal circuits and between hippocampal, amygdala, and hypothalamic regions, whereas cortical connectivity showed comparatively minor differences (Fig. 7J, left). Statistical comparison of the full co-activation matrices via a Mantel test revealed that there is a significant difference in overall network organization between male and female runners (Fig. 7J, left), indicating sex-dependent reorganization of network architecture following VWR. Permutation-based edge-level analyses identified multiple changes in female runners. Reduced correlations were observed between dGrDG and hippocampal (dCA1, dCA2, dCA3) and cortical (IL) regions, as well as between vGrDG and Pir, and between vPoDG and DLS (Fig. 7K, L). Additional reductions involved LHb connectivity with MO, VO, and LSD, as well as several hypothalamic and midbrain connections, including suprachiasmatic nucleus (SCN)–Arc, DMH–DLS, DMH–LO, PVN–LSV, VTA–LSV, and VTA–M2. Conversely, female runners exhibited increased correlations between amygdala and hippocampal regions (BMA–dCA2, BMA–dCA3) and between hypothalamic and hippocampal regions (LHA–dCA3, VMH–dCA3) relative to male runners (Fig. 7K, L).

Together, these findings indicate that VWR induces sex-specific reorganization of ΔFOSB co-activation patterns.

## DISCUSSION

Here, we demonstrate that long-term VWR in mice induced sustained neural activation in many brain regions associated with stress regulation, cognition and reward, as well as repression in a few brain regions. Network analyses revealed that VWR reorganized brain-wide ΔFOSB co-activation networks in a sex-specific manner compared to sedentary controls. Indeed, male runners showed marked topological reorganization towards a more cortico-central network architecture, accompanied by relatively modest changes in network density and global efficiency, whereas female runners exhibited fewer alterations in network topology, but a more pronounced increase in global efficiency.

### VWR-induced alterations in regional ΔFOSB expression

This study provides the first characterization of VWR-mediated ΔFOSB expression in the female mouse brain, revealing sustained activation of cortical regions. In addition, we identify novel VWR-mediated modulation of ΔFOSB in the male mouse brain, including in the lateral septum and PVT, that extends beyond previously identified modulation in striatal and hippocampal structures (Herrera et al., 2016; Mul et al., 2018; Nishijima et al., 2013).

Some of these modulations seem sex-specific. For example, VWR decreased PVT ΔFOSB in males, but trended to increase PVT ΔFOSB in females. To the best of our knowledge, a significant decrease in ΔFOSB, or a trend for a decrease, following VWR has to date been reported only for some brain regions: for the SCN in male and female running rats (Shiba et al., 2025), for the arcuate nucleus in female running rats (Hardonk et al., 2026), and for the basolateral amygdala in female running prairie voles (Watanasriyakul et al., 2019). Given the established role of the PVT in stress responsiveness, arousal and regulation of mesolimbic dopamine signaling (Elam et al., 2025; Ren et al., 2018), this finding may reflect a sex-specific recruitment of this stress-sensitive hub by long-term VWR. As chronic stress induces ΔFOSB accumulation in PVT neurons and suppression of these neurons produced antidepressant-like effects in male, but not female mice (Zhao et al., 2021), this suggests that a decreased ΔFOSB signaling in the male PVT may underlie the stress-buffering effects of VWR. However, because ΔFOSB function in the PVT remains poorly characterized, further studies are required to determine the functional significance of this modulation following VWR.

We also observed VWR-mediated suppression of ΔFOSB in the dGrDG of female mice. Whereas prior studies in female rats and prairie voles reported no VWR-induced modulation of dorsal hippocampal ΔFOSB (Hardonk et al., 2026; Watanasriyakul et al., 2019), the present findings suggest a sex-specific adaptation of the dentate gyrus to VWR. In male mice and rats, VWR seems to consistently increase dorsal hippocampal (DG) and ventral hippocampal (DG, CA1, CA3) ΔFOSB (Hardonk et al., 2026; Nishijima et al., 2013). In contrast, the present data indicate that long-term VWR results in reduced ΔFOSB accumulation in the female dGrDG. However, the opposing ΔFOSB changes between sexes may reflect sex-dependent routes towards an adaptive circuit configuration during long-term VWR, rather than opposing effects on hippocampal function. Future studies should assess the underlying mechanisms that drive the sex-dependent VWR modulation of dGrDG ΔFOSB and its subsequent functional implication.

Our mouse ΔFOSB following long-term VWR extends on our prior work in rats, and several patterns of VWR-induced ΔFOSB modulation were consistent between the current mouse study and our previously published rat ΔFOSB brain map following long-term VWR (Hardonk et al., 2026). For example, dorsal striatal, hippocampal and cingulate regions showed consistent induction in male rats and mice, whereas cortical and ventral hippocampal regions were predominantly affected in female rats and mice. These observations highlight core networks engaged by VWR in both species.

The number of brain regions that showed substantial VWR-mediated ΔFOSB induction, compared to sedentary controls, appeared lower in mice than in rats, and examples include medial prefrontal areas, AcbSh, multiple ventral dentate gyrus subregions and select amygdaloid and hypothalamic nuclei. Notably, ΔFOSB accumulation in the NAc, consistently reported in previous mouse and rat VWR studies (Greenwood et al., 2011; Herrera et al., 2016; Mul et al., 2018; Obici et al., 2015; Werme et al., 2002), was not observed in the current study. One important methodological difference is our use of light, low-resistance horizontal (“saucer-like”) plastic running wheels rather than the heavier, higher-resistance stainless-steel vertical wheels employed in most prior mouse and rat studies. Reduced physical demand may alter recruitment of effort- and reward-related circuits to sustain motivation for running, thereby possibly contributing to differences in ΔFOSB expression in limbic and motivational regions. Overall, these findings underscore both the conserved core networks engaged by VWR and the sensitivity of circuit recruitment to species and potentially running wheel properties, highlighting the need for direct comparisons across experimental conditions in future studies.

### VWR effects on ΔFOSB co-activation networks

As brain regions do not operate in isolationand ΔFOSB reflects patterns of sustained neural activity (McClung et al., 2004), we examined ΔFOSB co-activation networks. Previously, we found that long-term VWR in rats produced more efficient networks with reduced density and cortical-favored information flow, in both males and females (Hardonk et al., 2026). In the present study, VWR promoted a similar pattern in males, with an even stronger reorganization towards a cortex-centered architecture, accompanied by modest reductions in density and increased global efficiency. In females, VWR robustly increased global efficiency, consistent with prior rat data, but lacked density reduction or pronounced shifts towards cortical centrality. One possible explanation is that VWR induced the emergence of two distinct co-activation networks in female runners rather than a single integrated network. This may complicate the interpretation of local network characteristics to describe network centrality (*i.e.* node degree and betweenness centrality).

Notably, cortical regions constituted one of these networks, and applying more stringent thresholds (*P* < 0.0001) revealed that these within-cortex connections were stronger than the central nodes identified using a threshold of *P* < 0.005. An alternative, and not mutually exclusive, explanation is that sedentary females already display a more cortex-centered network organization at baseline, characterized by stronger and intracortical connectivity and lower overall network density. To test this, we directly compared ΔFOSB co-activation networks between sedentary males and females. Indeed, female sedentary animals showed higher cortical connections than sedentary males, together with fewer connections linking the dGrDG, amygdala and hypothalamic structures, as well as reduced connectivity involving the lateral habenula. After VWR, this female-specific cortical predominance was diminished, indicating that females start from a more cortex-centered, lower-density network configuration that may mask further VWR-induced cortical reorganization.

### Implications for stress resilience and future mechanistic studies

Our unique ΔFOSB brain map following long-term VWR offers a framework for future mechanistic studies into the impact of VWR-mediated changes in ΔFOSB in brain regions associated with stress regulation, cognition and reward. For example, prior studies in male mice demonstrated that stress- or VWR-induced ΔFOSB accumulation in the NAc (Mul et al., 2018; Vialou, Maze, et al., 2010; Vialou, Robison, et al., 2010), as well as stress induction of ΔFOSB in ventral hippocampus-to-NAc-projecting neurons, promoted stress resilience (Eagle et al., 2020), whereas stress-induced ΔFOSB in the PrL decreased stress resilience (Vialou et al., 2014). While we did not assess stress-related behavior in the present study, we identify several new regions previously not associated with VWR-mediated ΔFOSB modulation. These included the cingulate cortex and PVT in both sexes, the lateral septum in males and the orbital cortex in females. These findings highlight candidate regions for future experiments in which ΔFOSB signaling could be functionally manipulated to test their causal contribution in the stress resilience-promoting effects of long-term VWR.

In addition to identifying novel candidate brain regions modulated by VWR, our findings further revealed pronounced sex-specific differences in both regional ΔFOSB expression and brain-wide network organization. These sex differences may shape how exercise protects against chronic stress, which is relevant for the prevention of depression. Given the higher prevalence of depression in women (Ten Have et al., 2023) and the historical underrepresentation of females in mechanistic animal studies (Shansky, 2019), our findings highlight the importance of including sex as a biological variable in future mechanistic studies of VWR and its effects on stress regulation-related circuits.

## Conclusion

Taken together, we here demonstrate that long-term VWR in mice modulated ΔFOSB expression in brain regions associated with stress regulation-, cognition- and reward-related behavior and reorganized ΔFOSB co-activation networks with distinct sex-specific signatures. The current ΔFOSB brain atlas and changes in co-activation networks following VWR provide a framework for future functional studies in mice that may help elucidate more specific mechanistic pathways via which exercise training can promote brain health and stress resilience.

## Author contributions

M.H.H., J.D.M., S.E.l.F, P.J.L. designed research; M.H.H., R.W., J.S., J.G., J.D.M. performed research; A.H.V. contributed unpublished analytic tools; M.H.H., R.W., J.S., M.H.M. analyzed data; M.H.H., J.D.M., P.J.L. wrote the paper.

## Acknowledgements

This work was funded by Amsterdam Neuroscience (JDM) and the Centre for Urban Mental Health (JDM, and PJL). PJL is further supported by Alzheimer Nederland, by a ZonMw grant on biomedical research on ME/CFS (NMCB), by a ZonMW Memorabel grant on Alzheimer’s disease (MODEM) and is co-recipient of the NWO Gravitation grant ‘Institute for Chemical Neuroscience’ (iCNS; nr 024.006.009), funded by the Dutch Research Council (NWO). The authors would like to thank past and present members of the Fit Brain Lab (University of Amsterdam) and Brain Plasticity group (University of Amsterdam) for scientific feedback, insightful discussions, and their friendship. We especially thank Wouter Cox, Dorte Heideman, Daniël Marrinan, Gideon Meerhoff and Jan den Blaauwen for their contributions during various stages of the project. We also thank Rob de Heus and Gavin Adema for ensuring animal welfare. We thank the Netherlands Institute for Neuroscience microscopy support team Joris Coppens and Roeland Lokhorst for their expertise and assistance during image acquisition.

## Supplementary information

Supplementary information is available at the JoN’s website.

## Code availability

Access to the data and computational tools used in this study can be obtained from the corresponding author, upon reasonable request.

