## Supplementary information for "Sex-specific signatures of brain-wide induction of ΔFOSB and altered co-activation networks in a mouse model for exercise training"

### SUPPLEMENTAL FIGURES AND FIGURE LEGENDS

#### Supplementary figure 2-1

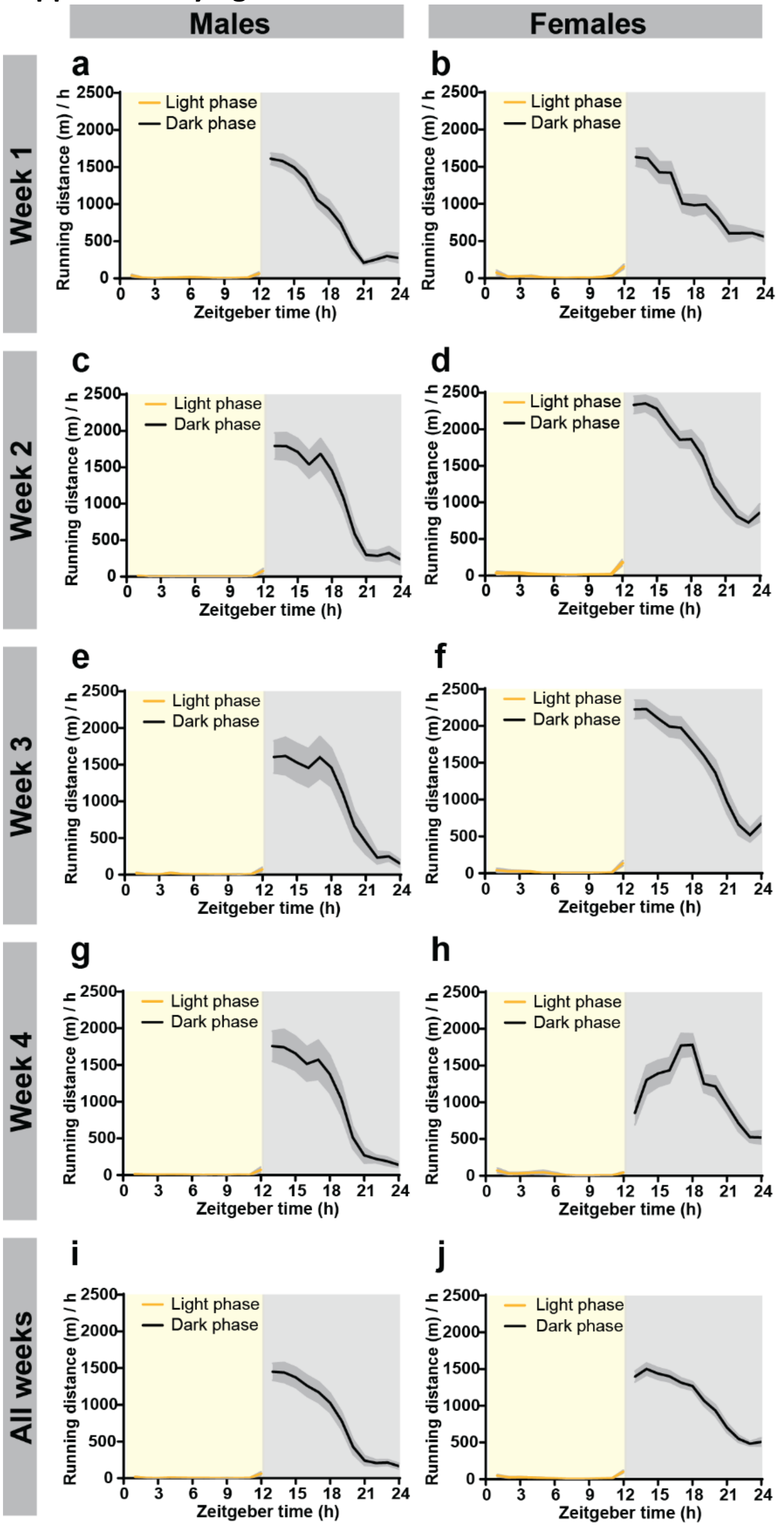


**Supplementary Figure 2-1. Circadian dynamics in daily VWR in males and females**. 24h running patterns averaged per running week (**a**-**h**) or averaged for all running weeks (**i**, **j**) in males (**a** ,**c**, **e**, **g**, **i**) and females (**b**, **d**, **f**, **h**, **j**). Zeitgeber Time (ZT) refers to time of day relative to the time of lights off/lights on, with lights off at ZT12. Data are presented as mean ± S.E.M. (a-j). a-j, *n* = 10-11/group.

**Supplementary figure 2-2**

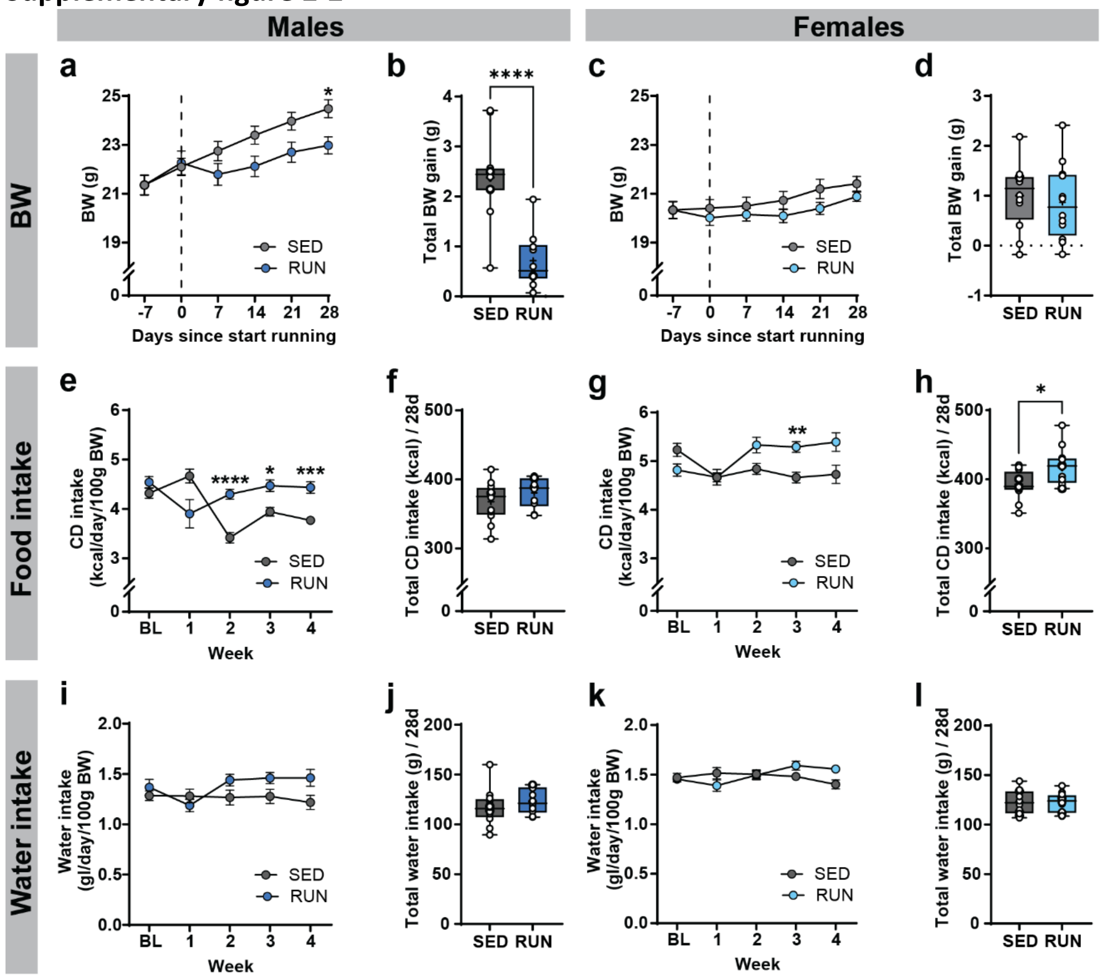


**Supplementary Figure 2-2. Impact of long-term VWR on body weight, food and water intake in males and females**. (**a**) Body weight (BW) evolution (*time x housing* interaction, $F_{(5, 100)}$ = 14.15, *P* < 0.0001; Šidák post hoc: **P* < 0.05, SED *versus* RUN) and (**b**) total BW gain ($t_{(20)}$ = 5.435, *****P* < 0.0001 *versus* SED) in male sedentary (SED) controls and runners (RUN). (**c**) BW evolution (*time x housing* interaction, $F_{(5, 110)}$ = 2.074, *P* = 0.074) and (**d**) total BW gain ($t_{(22)}$ = 0.469, *P* = 0.644 *versus* SED) in female SED and RUN mice. (**e**) Average daily control diet (CD) intake (*time x housing* interaction, $F_{(4, 80)}$ = 11.78, *P* < 0.0001; Šidák post hoc: *****P* < 0.0001, ****P* < 0.001, **P* < 0.05, SED *versus* RUN) and (**f**) total CD intake ($t_{(20)}$ = 1.180, *P* = 0.252 *versus* SED) in male SED and RUN mice. (**g**) Control diet (CD) intake evolution (*time x housing* interaction, $F_{(4, 88)}$ = 14.59, *****P* < 0.0001; Šidák post hoc: ***P* < 0.01, SED *versus* RUN) and (**h**) total CD intake ($t_{(22)}$ = 2.635, *P* = 0.015 *versus* SED) in female SED and RUN mice. (**i**) Average daily water intake (*time x housing* interaction, $F_{(4, 79)}$ = 8.390, *****P* < 0.0001) and (**j**) total water intake ($t_{(20)}$ = 0.932, *P* = 0.363 *versus* SED) in male SED and RUN mice. (**k**) Average daily water intake (*time x housing* interaction, $F_{(4, 88)}$ = 8.248, *****P* < 0.0001) and (**l**) total water intake ($t_{(22)}$ = 0.122, *P* = 0.904 *versus* SED) of female SED and RUN mice. Data are presented as the mean ± S.E.M (a, c, e, g, i, k) or as box plots indicating the median (line) and mean (+), the interquartile range, and the minimum to maximum values of the data distribution, with dots representing individual mice (SED = gray, RUN = blue; b, d, f, h, j, l). (a-l) *n* = 12/group.

Supplementary figure 2-3

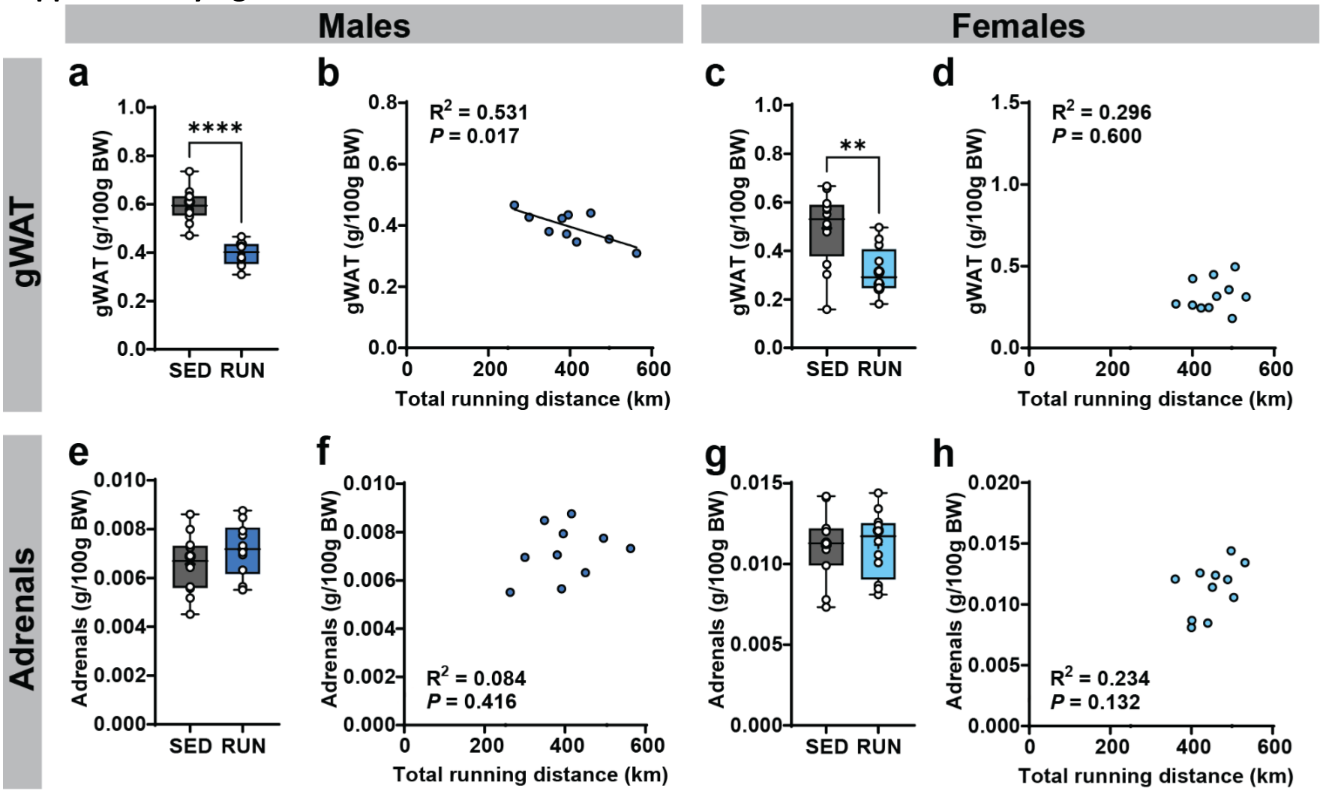


**Supplementary Figure 2-3. Physiological impact of VWR in males and females. (a)** Terminal gonadal white adipose tissue (gWAT) weight, normalized for body weight, in male sedentary (SED) controls and runners (RUN; $t_{(20)} = 7.652$, $P < 0.0001$ *versus* SED) and (**b**) correlation between terminal gWAT weight and total running distance ($R^2 = 0.531$, $F_{(1,8)} = 9.047$, $P = 0.017$) in male RUN mice. (**c**) Terminal gWAT weight, normalized for body weight, in female SED and RUN mice ($t_{(22)} = 3.384$, $P = 0.003$ *versus* SED) and (**d**) correlation between terminal gWAT weight and total running distance ($R^2 = 0.296$, $F_{(1,9)} = 0.396$, $P = 0.600$) in female RUN mice. (**e**) Terminal adrenal weight ($t_{(20)} = 1.222$, $P = 0.236$ *versus* SED) in male SED and RUN mice and (**f**) correlation between terminal adrenal weight and total running distance ($R^2 = 0.084$, $F_{(1,8)} = 0.737$, $P = 0.416$) in male RUN mice. (**g**) Terminal adrenals weight ($t_{(22)} = 0.070$, $P = 0.945$ *versus* SED) in female SED and RUN mice and (**h**) correlation between terminal adrenals weight and total running distance ($R^2 = 0.234$, $F_{(1,9)} = 2.742$, $P = 0.132$) in female RUN mice. Data are presented as box plots indicating the median (line) and mean (+), the interquartile range, and the minimum to maximum values of the data distribution, with dots representing individual mice (a, c, e, g) or as regression plots indicating the simple linear regression (line) with dots representing the individual mice (b, d, f, h). (a-h) $n$ = 10-12/group.

**Supplementary figure 3-1**

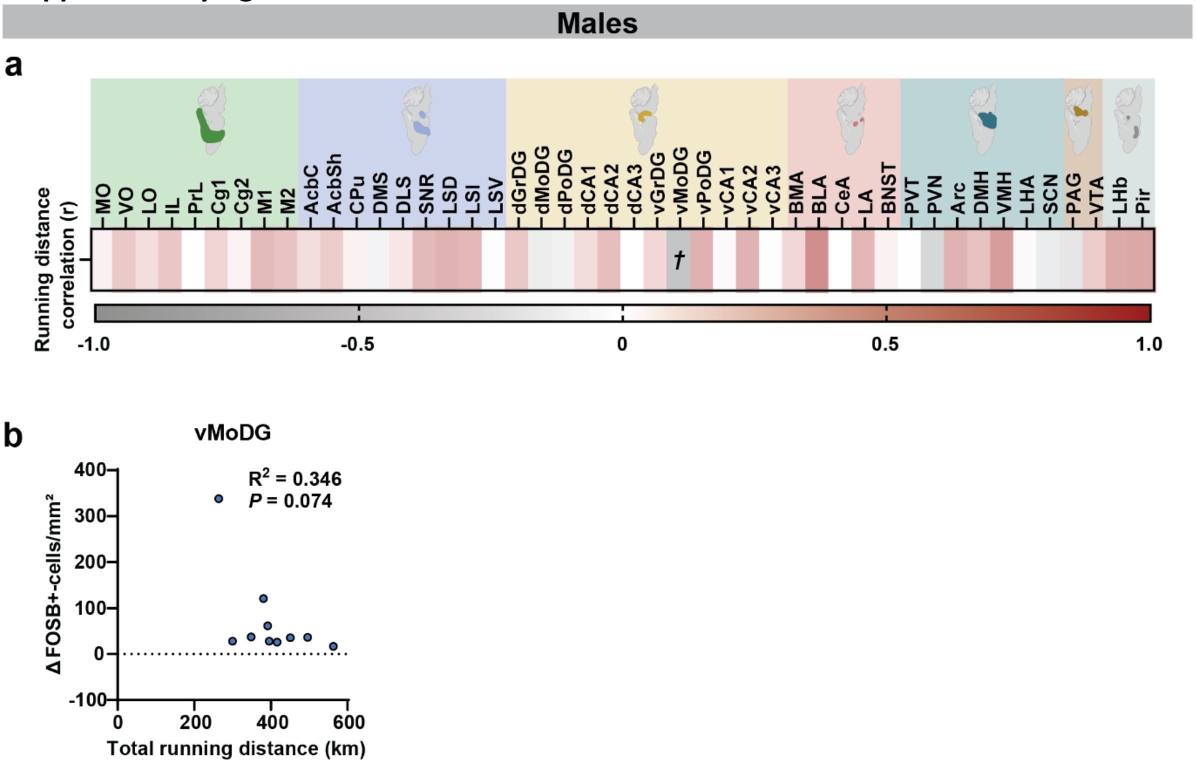


**Supplementary Figure 3-1. Pearson correlations between total running distance and brain ΔFOSB in males.** (**a**) Heatmap of Pearson correlation values (r) of total running distance and fold change in ΔFOSB-positive cells per region of interest (ROI) in male runners (RUN). See Supplementary Table 3-1 for the full names of the presented acronyms, †*P* < 0.10. (**b**) Correlation between total running distance and mean number of ΔFOSB-positive cells in (b) ventral molecular layer of the dentate gyrus (vMoDG; $R^2 = 0.346$, $F_{(1,8)} = 4.233$, $P = 0.074$) in male RUN mice. Data are presented as heatmaps indicating Pearson correlation coefficients (r) (a) or as regression plots indicating the simple linear regression (line) with dots representing individual mice (b). (a,b) *n* = 10/group.

**Supplementary figure 4-1**

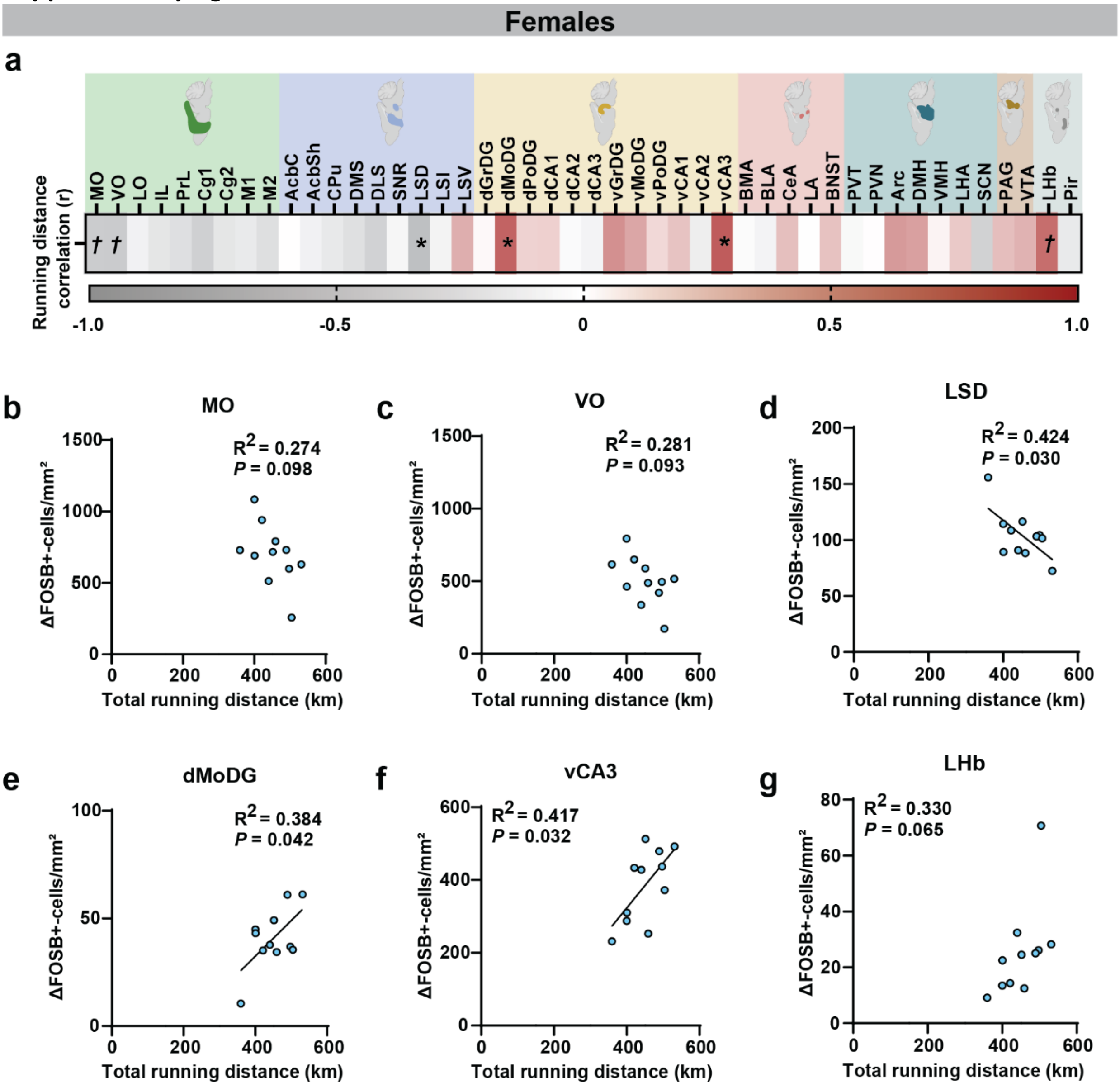


**Supplementary Figure 4-1. Pearson correlations between total running distance and brain ΔFOSB in females.** (**a**) Heatmap of Pearson correlations values (r) of total running distance and fold change in ΔFOSB-positive cells per region of interest (ROI) in female runners (RUN). See Supplementary Table 4-1 for the full names of the presented acronyms, **P* < 0.05, †*P* < 0.10. (**b-g**) Correlation between total running distance and mean number of ΔFOSB-positive cells in (b) medial (MO; $R^2 = 0.274$, $F_{(1,9)} = 3.403$, $P = 0.098$) and (c) ventral orbital cortex (VO; $R^2 = 0.281$, $F_{(1,9)} = 3.523$, $P = 0.093$), (d) dorsal part of the lateral septum (LSD; $R^2 = 0.424$, $F_{(1,9)}$ = 6;612, $P = 0.030$), (e) dorsal part of the molecular layer of the dentate gyrus (dMoDG; $R^2 = 0.384$, $F_{(1,9)} = 5.598$, $P = 0.042$), (f) ventral part of the cornus ammonis 3 (vCA3; $R^2 = 0.417$, $F_{(1,9)} = 6.427$, $P = 0.032$) and (g) lateral habenula (LHb; $R^2 = 0.330$, $F_{(1,9)} = 4.428$, $P = 0.065$). in female RUN mice. Data are presented as heatmaps indicating Pearson correlation coefficients (r) or as regression plots indicating the simple linear regression (line) with dots representing individual mice (b-g). (a-g) *n* = 11/group.

**Supplementary figure 5-1**

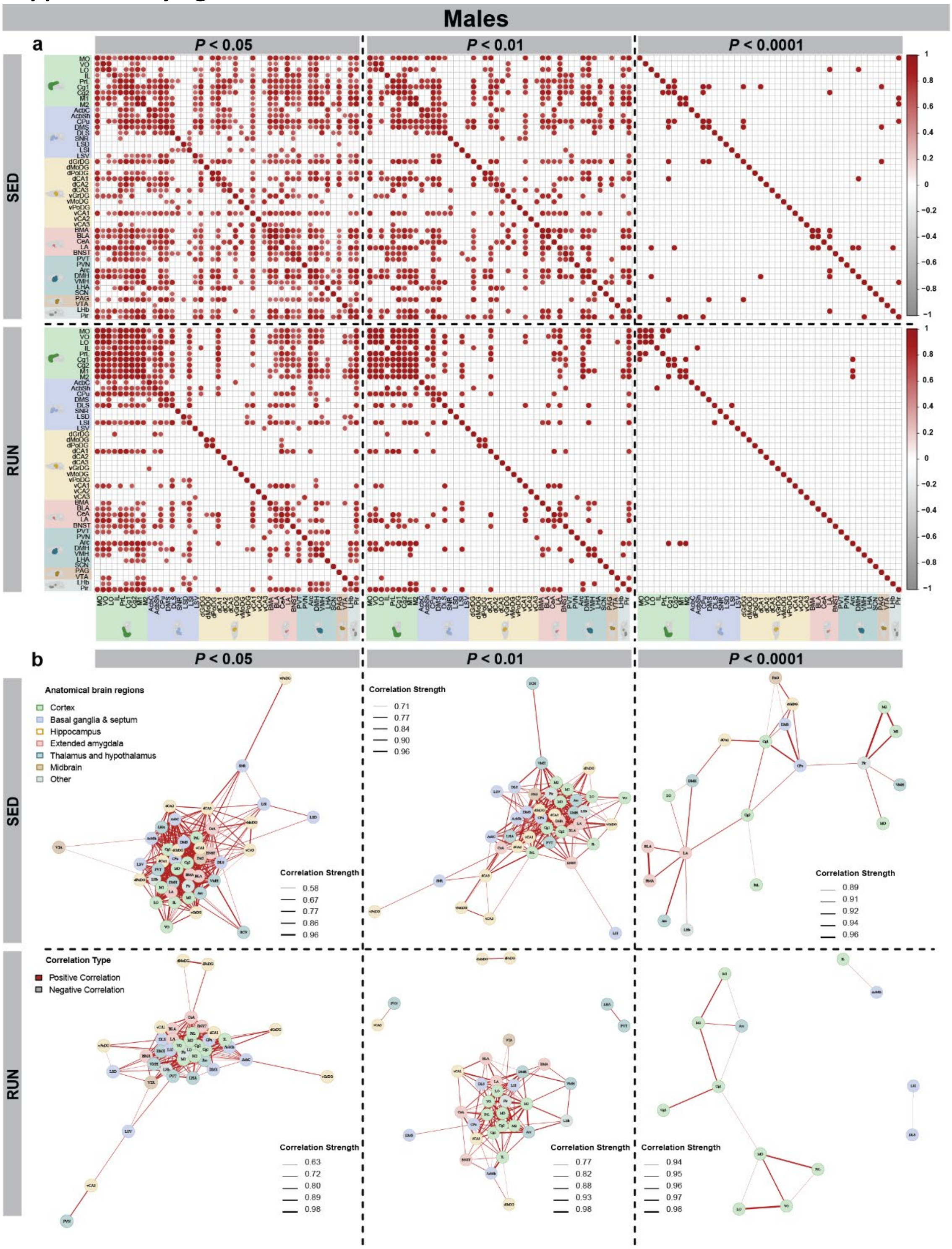


**Supplementary Fig. 5-1: The influence of thresholding on network topology in males.** (**a**) Regional cross-correlation matrices of ΔFOSB-positive cells between all pairs of neuroanatomical regions in male sedentary (SED) controls and runners (RUN) at thresholds of $\alpha < 0.05$, $\alpha < 0.01$ and $\alpha < 0.0001$. (**b**) Functional connectivity networks constructed of SED and RUN mice at thresholds of $\alpha < 0.05$, $\alpha < 0.01$ and $\alpha < 0.0001$. (a,b) $n$ = 10-12/group.

**Supplementary figure 6-1**

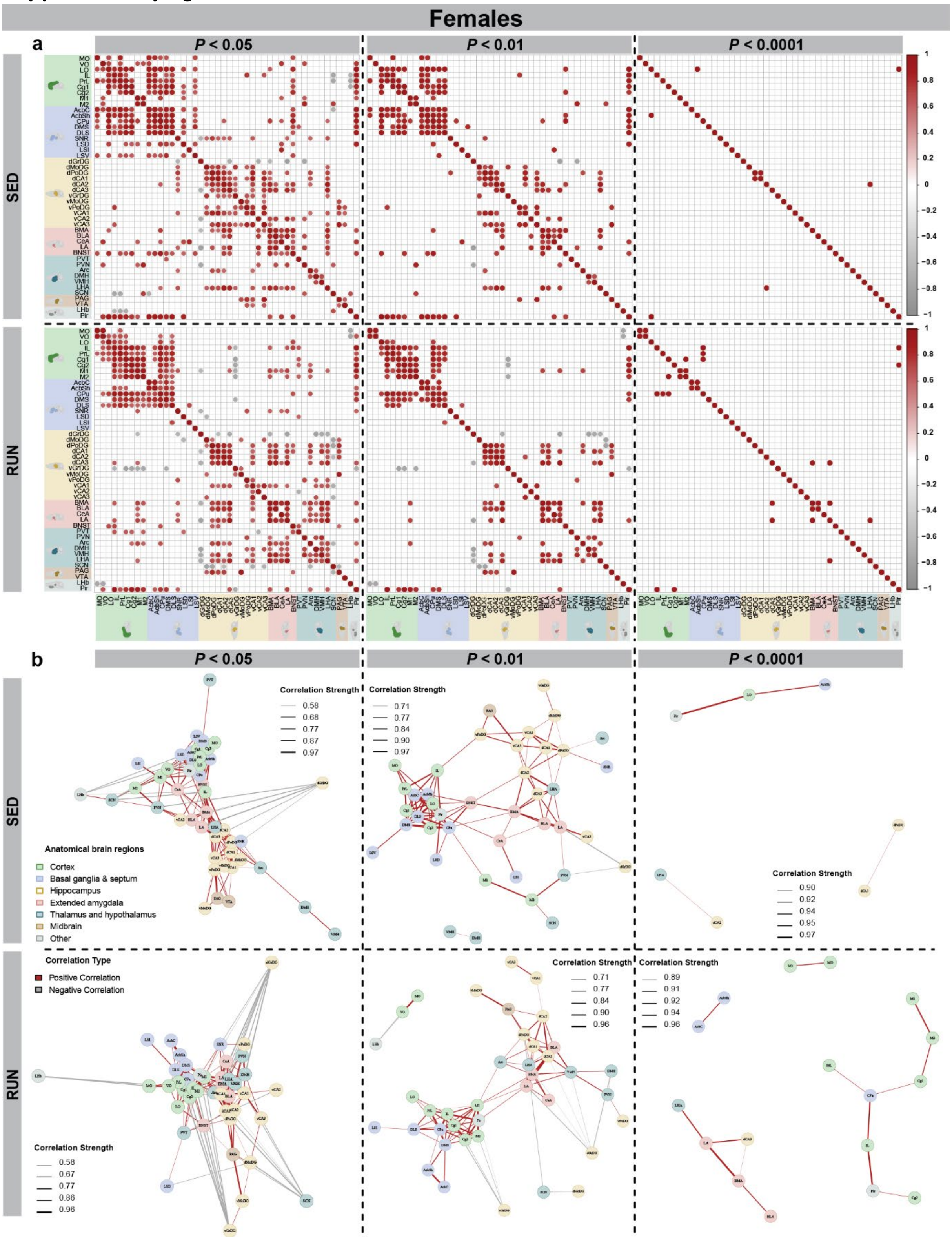


**Supplementary Fig. 6-1: The influence of thresholding on network topology in females.** (**a**) Regional cross-correlation matrices of ΔFOSB-positive cells between all pairs of neuroanatomical brain regions in female sedentary (SED) controls and runners (RUN) at thresholds of $\alpha < 0.05$, $\alpha < 0.01$ and $\alpha < 0.0001$. (**b**) Functional connectivity networks constructed of SED and RUN mice at thresholds of $\alpha < 0.05$, $\alpha < 0.01$ and $\alpha < 0.0001$. (a,b) $n = 12$/group.

### Supplementary table 3-1

| Brain region | Full name | *N* SED | *N* RUN | *% change* | *t/U* | *P* | *P* adjusted | Bregma range | *N* slices SED | *N* slices RUN |
|---|---|---|---|---|---|---|---|---|---|---|
| MO | *Medial orbital cortex* | 12 | 10 | -7.8% | 0.301 (t) | 0.767 | 0.855 | 2.80 – 1.98mm | 2 - 11 | 1 - 5 |
| VO | *Ventral orbital cortex* | 12 | 10 | 43.9% | -0.745 (t) | 0.467 | 0.766 | 2.80 – 1.98mm | 2 - 11 | 1 - 5 |
| LO | *Lateral orbital cortex* | 12 | 10 | 29.4% | -0.621 (t) | 0.542 | 0.779 | 2.80 – 1.70mm | 3 - 13 | 3 - 6 |
| IL | *Infralimbic cortex* | 12 | 10 | 7.1% | -0.265 (t) | 0.793 | 0.855 | 1.98 – 1.34mm | 2 - 5 | 1 - 4 |
| PrL | *Prelimbic cortex* | 12 | 10 | 6.9% | -0.275 (t) | 0.786 | 0.855 | 2.80 – 1.54mm | 4 - 13 | 3 - 6 |
| Cg1 | *Cingulate cortex 1* | 12 | 10 | 14.9% | -0.573 (t) | 0.573 | 0.779 | 2.34 – -0.22mm | 10 - 18 | 10 - 13 |
| Cg2 | *Cingulate cortex 2* | 12 | 10 | 54.8% | -2.065 (t) | 0.052 | 0.330 | 1.42 – -0.22mm | 6 - 9 | 6 - 8 |
| M1 | *Primary motor cortex* | 12 | 10 | 11.8% | -0.312 (t) | 0.759 | 0.855 | 2.34 – -1.34mm | 13 - 23 | 14 - 18 |
| M2 | *Supplementary motor cortex* | 12 | 10 | 28.5% | -0.758 (t) | 0.458 | 0.766 | 2.46 – -1.34mm | 13 - 24 | 14 - 18 |
| AcbC | *Nucleus accumbens core* | 12 | 10 | 12.1% | -0.715 (t) | 0.483 | 0.766 | 1.94 – 0.86mm | 5 - 6 | 4 - 5 |
| AcbSh | *Nucleus accumbens shell* | 12 | 10 | 4.2% | -0.209 (t) | 0.837 | 0.855 | 1.94 – 0.74mm | 5 - 6 | 4 - 5 |
| CPu | *Caudate putamen* | 12 | 10 | 25.4% | -1.363 (t) | 0.188 | 0.577 | 1.94 – -2.30mm | 17 - 20 | 16 - 19 |
| DMS | *Dorsomedial striatum* | 12 | 10 | 105.6% | -4.435 (t) | **0.001** | **0.013** | 1.54 – 0.14mm | 6 - 8 | 5 - 7 |
| DLS | *Dorsolateral striatum* | 12 | 10 | 87.2% | -2.094 (t) | 0.057 | 0.330 | 1.10 – 0.14mm | 4 - 7 | 4 - 6 |
| SNR | *Substantia nigra* | 12 | 10 | -57.3% | 1.573 (t) | 0.142 | 0.577 | -2.46 – -4.16mm | 4 - 8 | 5 - 7 |
| LSD | *Lateral septum, dorsal* | 12 | 10 | 40.5% | -1.446 (t) | 0.166 | 0.577 | 1.42 – -0.46mm | 6 - 9 | 6 - 9 |
| LSI | *Lateral septum, intermediate* | 12 | 10 | 81.4% | -26 (U) | **0.025** | 0.287 | 1.54 – -0.10mm | 6 - 7 | 5 - 7 |
| LSV | *Lateral septum, ventral* | 12 | 10 | 33.4% | -1.625 (t) | 0.121 | 0.559 | 1.18 – -0.10mm | 5 - 7 | 4 - 7 |
| dGrDG | *Granular layer of the dentate gyrus, dorsal* | 12 | 10 | 56.7% | -4.094 (t) | **0.001** | **0.013** | -0.94 – -3.64mm | 11 - 15 | 10 - 13 |
| dMoDG | *Molecular layer of the dentate gyrus, dorsal* | 12 | 10 | 12.5% | -0.684 (t) | 0.503 | 0.771 | -0.94 – -4.04mm | 11 - 16 | 11 - 15 |
| dPoDG | *Polymorph layer of the dentate gyrus, dorsal* | 12 | 10 | 23.7% | -0.838 (t) | 0.414 | 0.766 | -1.06 – -3.64mm | 11 - 14 | 9 - 13 |
| dCA1 | *Cornu ammonis 1, dorsal* | 12 | 10 | 33.7% | -1.618 (t) | 0.121 | 0.559 | -1.22 – -3.88mm | 10 - 15 | 10 - 14 |
| dCA2 | *Cornu ammonis 2, dorsal* | 12 | 10 | -8.1% | 0.358 (t) | 0.725 | 0.855 | -1.22 – -3.08mm | 6 - 10 | 6 - 9 |
| dCA3 | *Cornu ammonis 3, dorsal* | 12 | 10 | 3.3% | -0.212 (t) | 0.834 | 0.855 | -0.94 – -3.40mm | 8 - 12 | 8 - 11 |
| vGrDG | *Granular layer of the dentate gyrus, ventral* | 12 | 10 | 93.6% | -2.300 (t) | **0.034** | 0.298 | -2.46 – -3.88mm | 4 - 8 | 4 - 7 |
| vMoDG | *Molecular layer of the dentate gyrus, ventral* | 12 | 10 | 18.5% | -0.312 (t) | 0.759 | 0.855 | -2.46 – -4.04mm | 1 - 8 | 1 - 6 |
| vPoDG | *Polymorph layer of the dentate gyrus, ventral* | 12 | 10 | 59.6% | -1.373 (t) | 0.186 | 0.577 | -2.92 – --3.88mm | 2 - 8 | 3 - 5 |
| vCA1 | *Cornu ammonis 1, ventral* | 12 | 10 | 28.9% | -1.261 (t) | 0.222 | 0.637 | -2.80 – -3.88mm | 2 - 8 | 3 - 6 |
| vCA2 | *Cornu ammonis 2, ventral* | 10 | 7 | 75.5% | -2.351 (t) | **0.039** | 0.298 | -2.92 – -3.08mm | 1 - 2 | 1 - 2 |
| vCA3 | *Cornu ammonis 3, ventral* | 12 | 10 | 18.5% | -0.907 (t) | 0.375 | 0.766 | -2.80 – -3.64mm | 3 - 6 | 3 - 5 |
| BMA | *Basomedial amygdala* | 12 | 10 | -5.2% | 0.240 (t) | 0.813 | 0.855 | -0.58 – -2.70mm | 7 - 10 | 7 - 10 |
| BLA | *Basolateral amygdala* | 12 | 10 | 17.0% | -0.811 (t) | 0.429 | 0.766 | -0.58 – -3.16mm | 9 - 12 | 9 - 12 |
| CeA | *Central amygdala* | 12 | 10 | 19.5% | -1.014 (t) | 0.323 | 0.742 | -0.58 – -1.94mm | 4 - 8 | 5 - 7 |
| LA | *Lateral amygdala* | 12 | 10 | 13.4% | -0.507 (t) | 0.576 | 0.779 | -0.70 – -2.30mm | 5 - 8 | 5 - 8 |
| BNST | *Bed nucleus of the stria terminal* | 12 | 10 | 13.2% | -0.842 (t) | 0.410 | 0.766 | -0.62 – -2.18mm | 9 - 12 | 10 - 13 |
| PVT | *Paraventricular thalamic nucleus* | 12 | 10 | -45.1% | 2.849 (t) | **0.011** | 0.165 | -0.22 – -2.30mm | 2 - 6 | 3 - 6 |
| PVN | *Paraventricular hypothalamic nucleus* | 12 | 10 | -39.6% | 1.089 (t) | 0.291 | 0.706 | -0.58 – -1.22mm | 2 - 5 | 2 - 4 |
| Arc | *Arcuate nucleus* | 12 | 10 | -44.1% | 1.187 (t) | 0.253 | 0.684 | -1.22 – -2.80mm | 5 - 10 | 5 - 8 |
| DMH | *Dorsomedial hypothalamus* | 12 | 10 | 50.8% | -1.475 (t) | 0.159 | 0.577 | -1.46 – -2.18mm | 3 - 6 | 2 - 5 |
| VMH | *Ventromedial hypothalamus* | 12 | 10 | -2.5% | 0.052 (t) | 0.959 | 0.959 | -1.06 – -2.06mm | 3 - 7 | 3 - 5 |
| LHA | *Lateral hypothalamus* | 12 | 10 | 13.6% | -0.732 (t) | 0.474 | 0.766 | -0.34 – -2.80mm | 8 - 12 | 9 - 12 |
| SCN | *Suprachiasmatic nucleus* | 12 | 10 | -41.3% | 0.973 (t) | 0.343 | 0.751 | -0.22 – -0.82mm | 1 - 5 | 3 - 4 |
| PAG | *Periaqueductal grey* | 12 | 10 | 11.0% | -0.592 (t) | 0.562 | 0.779 | -2.70 – -3.28mm | 4 - 7 | 5 - 8 |
| VTA | *Ventral tegmental area* | 12 | 10 | -37.9% | 1.132(t) | 0.274 | 0.700 | -2.92 – -3.88mm | 2 - 5 | 3 - 5 |
| LHb | *Lateral habenula* | 12 | 10 | -15.4% | 0.426 (t) | 0.674 | 0.855 | -0.94 – -1.94mm | 3 - 7 | 4 - 5 |
| Pir | *Piriform cortex* | 12 | 10 | 7.7% | -0.308 (t) | 0.761 | 0.855 | 2.46 – -2.80mm | 19 - 28 | 20 - 24 |

**Supplementary Table 3-1**: **Quantification of ΔFOSB per brain region of interest in males**. Fold change in ΔFOSB-positive cells between male sedentary (SED) controls and runners (RUN). Data were compared with an independent Student's t-test (indicated with *t*) or a Mann-Whitney U-test (indicated with *U*) depending on data distribution. *N SED/N RUN* = number of mice; *P* = nominal *p*-value; *P* adjusted = nominal *P*-values after FDR correction for multiple comparisons; *% change* = fold change in ΔFOSB-positive cells in RUN mice compared to SED controls; Bregma range = Bregma ranges included per brain region; *N* slices = range of the min and max number of slices per brain region per group. Significant *P*-values (< 0.05) are marked in bold.

### Supplementary table 4-1

| Brain region | Full name | *N* SED | *N* RUN | *% change* | *t/U* | *P* | *P* adjusted | Bregma range | *N* slices SED | *N* slices RUN |
|---|---|---|---|---|---|---|---|---|---|---|
| MO | *Medial orbital cortex* | 9 | 12 | 40% | -2.427 (t) | **0.025** | 0.389 | 2.80 – 1.98mm | 1 - 4 | 1 - 4 |
| VO | *Ventral orbital cortex* | 9 | 12 | 30.2% | -1.829 (t) | 0.083 | 0.427 | 2.80 – 1.98mm | 1 - 4 | 1 - 4 |
| LO | *Lateral orbital cortex* | 10 | 12 | 13.1% | -0.683 (t) | 0.503 | 0.765 | 2.80 – 1.70mm | 1 - 5 | 2 - 5 |
| IL | *Infralimbic cortex* | 10 | 11 | 31.0% | -1.526 (t) | 0.144 | 0.530 | 1.98 – 1.34mm | 1 - 4 | 2 - 5 |
| PrL | *Prelimbic cortex* | 10 | 12 | 29.6% | -2.227 (t) | **0.038** | 0.427 | 2.80 – 1.54mm | 1 - 6 | 2 - 7 |
| Cg1 | *Cingulate cortex 1* | 10 | 12 | 32.9% | -31 (U) | 0.059 | 0.427 | 2.34 – -0.22mm | 4 - 14 | 6 - 14 |
| Cg2 | *Cingulate cortex 2* | 10 | 12 | 63.3% | -4.245 (t) | **0.001** | **0.021** | 1.42 – -0.22mm | 4 - 9 | 4 - 10 |
| M1 | *Primary motor cortex* | 12 | 12 | 28.6% | -50 (U) | 0.219 | 0.530 | 2.34 – -1.34mm | 2 - 18 | 9 - 21 |
| M2 | *Supplementary motor cortex* | 12 | 12 | 27.8% | -48 (U) | 0.178 | 0.530 | 2.46 – -1.34mm | 2 - 18 | 9 - 22 |
| AcbC | *Nucleus accumbens core* | 10 | 12 | 7.2% | -48 (U) | 0.456 | 0.765 | 1.94 – 0.86mm | 4 - 6 | 1 - 6 |
| AcbSh | *Nucleus accumbens shell* | 10 | 12 | 3.7% | -54 (U) | 0.722 | 0.831 | 1.94 – 0.74mm | 1 - 6 | 1 - 6 |
| CPu | *Caudate putamen* | 12 | 12 | 13.1% | -63 (U) | 0.630 | 0.801 | 1.94 – -2.30mm | 8 - 21 | 13 - 26 |
| DMS | *Dorsomedial striatum* | 10 | 12 | 32.4% | -32 (U) | 0.069 | 0.427 | 1.54 – 0.14mm | 4 - 7 | 2 - 8 |
| DLS | *Dorsolateral striatum* | 10 | 12 | -5.1% | -73 (U) | 0.418 | 0.765 | 1.10 – 0.14mm | 3 - 6 | 1 - 8 |
| SNR | *Substantia nigra* | 12 | 12 | -0.1% | 0.009 (t) | 0.998 | 0.998 | -2.46 – -4.16mm | 4 - 10 | 4 - 10 |
| LSD | *Lateral septum, dorsal* | 11 | 12 | 15.7% | -57 (U) | 0.608 | 0.801 | 1.42 – -0.46mm | 1 - 10 | 5 - 13 |
| LSI | *Lateral septum, intermediate* | 10 | 12 | 12.0% | -0.322 (t) | 0.751 | 0.834 | 1.54 – -0.10mm | 4 - 8 | 3 - 9 |
| LSV | *Lateral septum, ventral* | 10 | 12 | 7.7% | -0.473 (t) | 0.642 | 0.801 | 1.18 – -0.10mm | 5 - 8 | 3 - 9 |
| dGrDG | *Granular layer of the dentate gyrus, dorsal* | 12 | 12 | -42.0% | 3.930 (t) | **0.001** | **0.021** | -0.94 – -3.64mm | 12 - 16 | 10 - 24 |
| dMoDG | *Molecular layer of the dentate gyrus, dorsal* | 12 | 12 | 11.6% | -0.839 (t) | 0.411 | 0.765 | -0.94 – -4.04mm | 11 - 17 | 11 - 24 |
| dPoDG | *Polymorph layer of the dentate gyrus, dorsal* | 12 | 12 | -8.5% | 0.667 (t) | 0.512 | 0.765 | -1.06 – -3.64mm | 9 - 14 | 10 - 20 |
| dCA1 | *Cornu ammonis 1, dorsal* | 12 | 12 | -0.3% | 0.038 (t) | 0.970 | 0.991 | -1.22 – -3.88mm | 10 - 18 | 10 - 23 |
| dCA2 | *Cornu ammonis 2, dorsal* | 12 | 12 | 18.1% | -1.290 (t) | 0.211 | 0.530 | -1.22 – -3.08mm | 7 - 11 | 7 - 16 |
| dCA3 | *Cornu ammonis 3, dorsal* | 12 | 12 | 7.9% | -0.649 (t) | 0.523 | 0.765 | -0.94 – -3.40mm | 9 - 13 | 8 - 18 |
| vGrDG | *Granular layer of the dentate gyrus, ventral* | 12 | 12 | 24.0% | -1.153 (t) | 0.261 | 0.597 | -2.46 – -3.88mm | 4 - 10 | 4 - 12 |
| vMoDG | *Molecular layer of the dentate gyrus, ventral* | 12 | 12 | -12.9% | 0.494 (t) | 0.627 | 0.801 | -2.46 – -4.04mm | 1 - 8 | 2 - 7 |
| vPoDG | *Polymorph layer of the dentate gyrus, ventral* | 12 | 12 | 64.9% | -1.829 (t) | 0.083 | 0.427 | -2.92 – --3.88mm | 2 - 8 | 2 - 8 |
| vCA1 | *Cornu ammonis 1, ventral* | 12 | 12 | 24.6% | -1.467 (t) | 0.159 | 0.530 | -2.80 – -3.88mm | 3 - 9 | 2 - 9 |
| vCA2 | *Cornu ammonis 2, ventral* | 11 | 11 | -13.4% | 0.473 (t) | 0.644 | 0.801 | -2.92 – -3.08mm | 1 - 2 | 1 - 3 |
| vCA3 | *Cornu ammonis 3, ventral* | 12 | 12 | 18.0% | -1.441 (t) | 0.164 | 0.530 | -2.80 – -3.64mm | 4 - 6 | 3 - 10 |
| BMA | *Basomedial amygdala* | 12 | 12 | 14.1% | -0.929 (t) | 0.365 | 0.731 | -0.58 – -2.70mm | 6 - 11 | 7 - 12 |
| BLA | *Basolateral amygdala* | 12 | 12 | 8.0% | -0.635 (t) | 0.532 | 0.765 | -0.58 – -3.16mm | 8 - 13 | 11 - 20 |
| CeA | *Central amygdala* | 12 | 12 | 15.9% | -58 (U) | 0.443 | 0.765 | -0.58 – -1.94mm | 6 - 9 | 6 - 11 |
| LA | *Lateral amygdala* | 12 | 12 | 10.2% | -0.718 (t) | 0.480 | 0.765 | -0.70 – -2.30mm | 5 - 9 | 6 - 13 |
| BNST | *Bed nucleus of the stria terminal* | 12 | 12 | 13.0% | -55 (U) | 0.347 | 0.726 | -0.62 – -2.18mm | 7 - 14 | 9 - 18 |
| PVT | *Paraventricular thalamic nucleus* | 12 | 12 | 40.2% | -1.830 (t) | 0.083 | 0.427 | -0.22 – -2.30mm | 2 - 5 | 3 - 6 |
| PVN | *Paraventricular hypothalamic nucleus* | 12 | 12 | 48.2% | -1.138 (t) | 0.273 | 0.597 | -0.58 – -1.22mm | 1 - 4 | 1 - 6 |
| Arc | *Arcuate nucleus* | 12 | 12 | 7.4% | -0.222 (t) | 0.827 | 0.884 | -1.22 – -2.80mm | 5 - 9 | 3 - 14 |
| DMH | *Dorsomedial hypothalamus* | 12 | 12 | 5.5% | -0.308 (t) | 0.761 | 0.834 | -1.46 – -2.18mm | 4 - 6 | 3 - 8 |
| VMH | *Ventromedial hypothalamus* | 12 | 12 | 24.9% | -65 (U) | 0.713 | 0.831 | -1.06 – -2.06mm | 3 - 6 | 3 - 8 |
| LHA | *Lateral hypothalamus* | 12 | 12 | 15.8% | -48 (U) | 0.178 | 0.530 | -0.34 – -2.80mm | 9 - 13 | 8 - 18 |
| SCN | *Suprachiasmatic nucleus* | 12 | 12 | -52.4% | 1.460 (t) | 0.168 | 0.530 | -0.22 – -0.82mm | 1 - 4 | 3 - 6 |
| PAG | *Periaqueductal grey* | 12 | 12 | 18.8% | -1.299 (t) | 0.209 | 0.530 | -2.70 – -3.28mm | 2 - 8 | 3 - 11 |
| VTA | *Ventral tegmental area* | 12 | 12 | -9.2% | 0.388 (t) | 0.702 | 0.831 | -2.92 – -3.88mm | 2 - 7 | 2 - 7 |
| LHb | *Lateral habenula* | 12 | 12 | -1.4% | 0.062 (t) | 0.951 | 0.991 | -0.94 – -1.94mm | 5 - 6 | 4 - 10 |
| Pir | *Piriform cortex* | 12 | 12 | 10.9% | -1.270 (t) | 0.218 | 0.530 | 2.46 – -2.80mm | 8 - 26 | 13 - 26 |

**Supplementary Table 4-1**: **Quantification of ΔFOSB per brain region of interest in females**. Fold change in ΔFOSB-positive cells between female sedentary (SED) controls and runners (RUN). Data were compared with an independent Student's t-test (indicated with *t*) or a Mann-Whitney U-test (indicated with *U*) depending on data distribution. *N SED/N RUN* = number of mice; *P* = nominal *p*-value; *P* adjusted = nominal *P*-values after FDR correction for multiple comparisons; *% change* = fold change in ΔFOSB-positive cells in RUN mice compared to SED controls; Bregma range = Bregma ranges included per brain region; *N* slices = range of the min and max number of slices per brain region per group. Significant *P*-values (< 0.05) are marked in bold.
